# The phylogenetic context for the origin of a unique purple-green photosymbiosis

**DOI:** 10.64898/2025.12.20.694765

**Authors:** Sergio A. Muñoz-Gómez, Megan E.S. Sørensen, Shahed U.A. Shazib, Mann Kyoon Shin, Thilo Bauer, Martin Kreutz, Sebastian Hess

## Abstract

Symbioses are widespread in nature and have been the source of much evolutionary innovation. While some types of symbioses evolved multiple times across space and time (e.g., oxygenic photosymbioses or chemosymbioses), others are extremely rare. Purple photosymbioses are an example of such rare associations. Only two purple photosymbioses between heterotrophic eukaryotes and intracellular purple bacteria have been documented. This is in stark contrast to very common oxygenic photosymbioses and poses the question of what factors prevent the more frequent establishment of purple photosymbioses. To shed light on this question, we investigated the evolutionary history of the purple-green ciliate *Pseudoblepharisma tenue* (Spirostomidae) using a phylogenetic and comparative approach. We sampled about 30 new isolates of spirostomid ciliates, inferred a comprehensive and well-supported phylogenomic tree, and resolved the sister relationship between *Pseudoblepharisma* and *Spirostomum*. Furthermore, we characterized *P. tenue*’s sister species, here renamed *Pseudoblepharisma chlorelligera*, and revealed that it constitutes a quadripartite symbiosis between a ciliate, a green alga, and two different non-photosynthetic bacteria. In addition, we discovered three colorless, non-photosymbiotic *Pseudoblepharisma* species, which branch as sister to the photosymbiotic *P. tenue* and *P. chlorelligera*. Our phylogenetic and comparative genomic analyses suggest that the green algal symbionts of *P. tenue* predated the acquisition of purple bacterial symbionts, and that the ancestor of the extant *Pseudoblepharisma* species was non-photosymbiotic and facultatively anaerobic. These data allowed us to hypothesize on the evolutionary steps that led to the origin of *P. tenue* and thus bring us closer to explaining the conditions that led to the evolutionary emergence of a unique purple-green symbiosis.

## Introduction

Symbioses between heterotrophic eukaryotes and intracellular purple bacteria—purple photosymbioses—are extraordinarily uncommon in nature [1]. Only two examples have ever been reported, the oligotrich ciliate *Strombidium purpureum* and the heterotrich ciliate *Pseudoblepharisma tenue* [2, 3]. Even though both ciliates harbor anoxygenic photosynthesizers (purple bacteria) as intracellular symbionts, they differ considerably in physiology and ecology [1, 4, 5]. In contrast to these purple photosymbioses, classical photosymbioses with oxygenic photosynthesizers (eukaryotic algae or cyanobacteria) are phylogenetically and environmentally widespread [6]. Similarly, chemosymbioses comprising eukaryotic hosts and chemosynthetic (autotrophic) bacteria occur across diverse marine environments (e.g., shallow-water seagrass beds and the deep sea) and have evolved multiple times [7]. It remains unclear why purple photosymbioses, in particular, are so rare in nature.

*P. tenue* is a unique photosymbiosis as it represents the only eukaryote known to contain two contrasting photosynthetic endosymbionts: oxygenic green algae and anaerobic purple bacteria (**Fig. 1A, 1B**) [3, 8, 9]. Phylogenetic analyses with commonly used marker genes revealed that the ciliate host is most closely related to the genus *Spirostomum*, while the green algal symbionts belong to the genus *Chlorella*. The purple bacterial symbionts were identified as members of the *Chromatiaceae* (*Gammaproteobacteria*) and described as “*Candidatus* Thiodictyon intracellulare”. Interestingly, the purple bacteria occupy a volume 20 times larger than that of the green algal symbionts in the host cell (**Fig. 1A**). This strongly suggests that the physiology of the symbiotic consortium is dictated by the metabolic activity of the purple bacterial symbionts. Furthermore, “*Ca.* Thiodictyon intracellulare” experienced considerable genomic reduction and physiological specialization compared to their closest free-living relatives. For example, these purple bacterial symbionts lost the capacity to fix nitrogen and use sulfur compounds as photosynthetic electron donors. The provision of photosynthetically fixed carbon to the host—using molecular hydrogen or oxidized carbon sources (e.g., acetate) as electron donors—might be the primary contribution of “*Ca.* Thiodictyon intracellulare” to the symbiosis.

**Figure 1.**
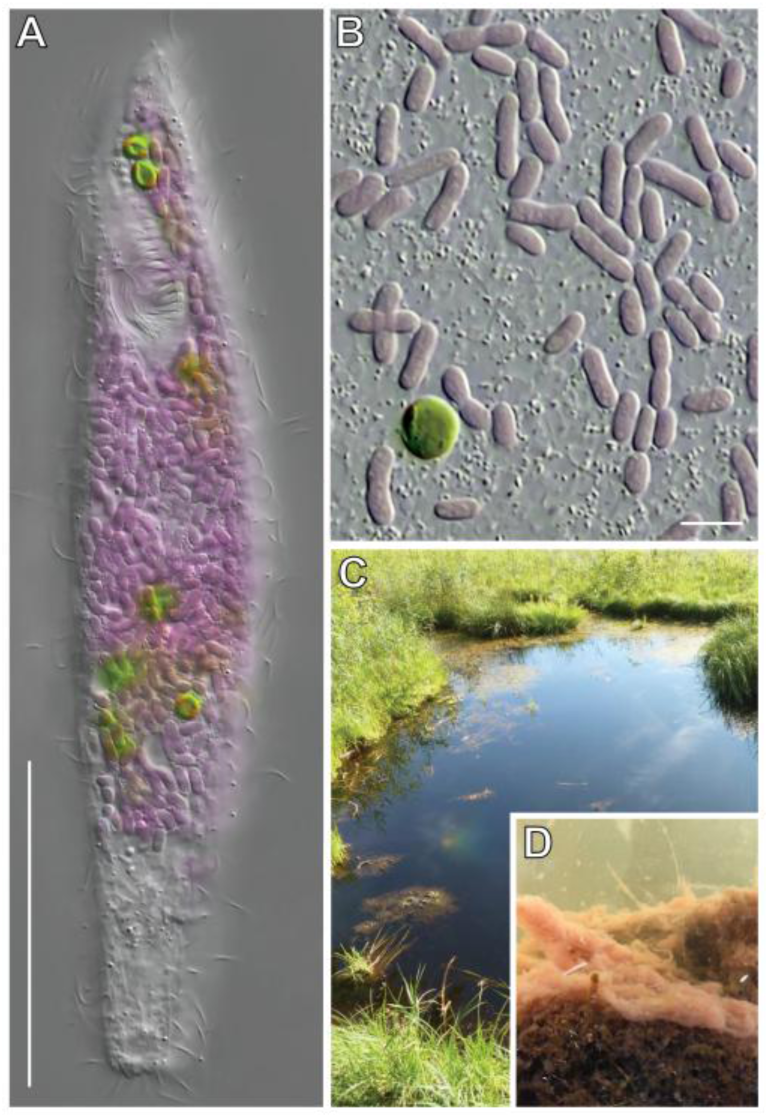
*P. tenue* and its two contrasting photosynthetic endosymbionts. A. The heterotrich ciliate *P. tenue*. B. Purple bacterial symbionts and a green algal symbiont cell released from the cytoplasm of *P. tenue*. C. Simmelried ponds where *P. tenue* is consistently found throughout the year. D. Freshwater sediments from Simmelried ponds that *P. tenue* inhabits. These sediments are rich in organic matter and occasionally exhibit outgrowths of purple sulfur bacteria. Scale bars: 50 µm (A), 5 µm (B).

*P. tenue* inhabits organic matter-rich and oxygen-poor sediments in relatively shallow freshwater ponds (**Fig. 1C, 1D**) [3, 10]—these same sediments often support the growth of various purple sulfur bacteria (see Fig. S1 in [3]). Metabolic reconstructions suggested that the *P. tenue* symbiotic consortium is physiologically flexible as each symbiotic partner has two or more modes of energy conservation [3]. We previously hypothesized on the major metabolic exchanges that sustain *P. tenue* under different environmental conditions [3]. In light and hypoxia, the purple symbionts might derive electrons from fermentation end-products of the ciliate host and/or green algal symbionts (e.g., hydrogen gas and acetate), and provide reduced photosynthate in return. In dark and oxia, each symbiotic partner switches to (micro)aerobic respiration to oxidize photosynthate reserves or phagocytosed food.

We currently do not know how two contrasting photosynthetic symbionts came to exist within the same host, the order in which the intracellular symbionts were acquired, and what pre-adaptations facilitated symbiont acquisition. Moreover, the precise phylogenetic placement of *Pseudoblepharisma* among spirostomid ciliates—a prerequisite to understand the evolution of its endosymbiosis—remains unclear [3, 11]. Progress on these questions is hampered by the fact that species closely related to *P. tenue* are unknown or lack genetic data for their endosymbionts. A purely green *Chlorella*-harboring spirostomid ciliate, originally described as *P. tenue* var. *chlorelligera* [12, 13] and recently referred to as *P. tenue* var. *viride* [10, 14], has been documented from European and North American freshwater ecosystems. A population of this purely green ciliate from Florida (U.S.A.) was recently shown to be phylogenetically most closely related to the purple-green *P. tenue* [14]. However, the limited resolution of the 18S rRNA gene and the lack of closely related taxa in these analyses prevents any conclusive phylogenetic interpretation.

Here, we took a phylogenetic and comparative approach to understand the evolutionary history of *P. tenue* and its close relatives. We sampled >30 populations of *Spirostomum*-like ciliates from diverse environments and sequenced single-cell amplified genomes and/or transcriptomes from the most phylogenetically disparate representatives. We inferred the most comprehensive and well-supported multiprotein tree of the family Spirostomidae to date using state-of-the-art phylogenomic analyses. These analyses revealed that four of the sampled populations constitute species that are more closely related to *P. tenue* than to any other species in the family Spirostomidae. One of these species corresponds to the above-mentioned purely green ciliate, here formally renamed *Pseudoblepharisma chlorelligera stat. nov*. We report on the morphology, ultrastructure, metagenome, and inferred metabolism of this species and uncover the quadripartite nature of its symbiosis which comprises a green algal and two bacterial endosymbionts. We furthermore present three colorless *Pseudoblepharisma* species that together form a sister clade to the photosymbiotic *Pseudoblepharisma* species (*P. tenue* and *P. chlorelligera*) and discuss the implications of our findings for understanding the evolutionary history of the unique purple-green symbiosis of *P. tenue*.

## Results and Discussion

### Expanding the known diversity of the family Spirostomidae reveals close relatives of *Pseudoblepharisma tenue*

To find potentially new populations that may be closely related to *P. tenue* and thus shed light on its evolutionary history, we sampled diverse locations across Germany, Austria, and South Korea in search for *Pseudoblepharisma*- and *Spirostomum*-like cells (**Table S1**). These cells were either directly isolated from environmental samples or enrichment cultures. Most of them were colorless and heterotrophic, except for green alga-harboring *Spirostomum semivirescens* [15] and the aforementioned purely green *Pseudoblepharisma*-like cell that has been referred to as *P. tenue* var. *chlorelligera* [12, 13] or *P. tenue* var. *viride* [10, 14].

The phylogeny of the Spirostomidae based on the rRNA gene operon and the mitochondrial COI gene shows that many of the newly isolated cells represent new strains or potential species (**Fig. S1**). This analysis confirms that *P. tenue* var. *chlorelligera* (Simmelried strain) is genetically distinct and branches as a sister to *P. tenue* (**Fig. S1**), warranting a change in taxonomic status and hereafter referred to as *P. chlorelligera* (see below for additional supporting evidence). (Whether the Simmelried and Florida strains are conspecific remains uncertain due to several ambiguous sites in the 18S rRNA gene sequence of the Florida strain.) Interestingly, some of the new heterotrophic spirostomids appear to be most closely related to photosymbiotic *Pseudoblepharisma* species or be sister to the entire *Spirostomum*‒*Pseudoblepharisma* clade (**Fig. S1**). The support for the phylogenetic placement of the genus *Pseudoblepharisma* and the new isolates, however, remained only low or moderate in this oligo-gene phylogeny (**Fig. S1**) [3, 14].

To robustly resolve the phylogeny of the family Spirostomidae, we chose the most phylogenetically representative strains and isolates (based on **Fig. S1**) for single-cell whole genome or transcriptome amplification and deep short-read sequencing (see **Table S2**). We then inferred comprehensive phylogenetic trees based on 203 proteins using site-heterogeneous models of protein sequence evolution, including an expanded set of recently sequenced *Spirostomum* strains [16, 11]. The resulting phylogeny is strongly supported in all of its basal branches, confirms the sisterhood of *P. tenue* and *P. chlorelligera* (Simmelried strain), and reveals that three heterotrophic isolates, namely TBCC008, TBCC048, and PsK1, form the sister clade of the photosymbiotic *Pseudoblepharisma* species (**Fig. 2A-C, S2, S3, S4**). The three heterotrophic isolates (**Fig. 2D-F**) came from freshwater sediments rich in decaying organic matter. Isolate PsK1 (**Fig. 2D, S5**), which was found in Ulsan, South Korea, displayed an elliptic macronucleus. Its closest relative, isolate TBCC048 (**Fig. 2E**) from Regenstauf, Germany, also possessed an elliptic macronucleus, which is similar to what is observed in the photosymbiotic species *P. tenue* and *P. chlorelligera*. Furthermore, isolate TBCC048 superficially resembled *Pseudoblepharisma crassum* KAHL 1927, which was described as a heterotroph preferentially feeding on purple bacteria (“rhodobakterien”) nearly hundred years ago [17]. On the other hand, isolate TBCC008 from Rheinbach, Germany, had a moniliform macronucleus with 6-8 nodules and might represent a new species (to be described in more detail elsewhere) (**Fig. 2F**). The genus *Pseudoblepharisma* is thus composed of two subgroups: (1) a photosymbiotic clade of species that carry at least green algal endosymbionts, and (2) a purely heterotrophic clade (**Fig. 2A**). The *Pseudoblepharisma* clade confidently branches as sister to the *Spirostomum* clade (**Fig. 2A**); the support for this branch increased as more complex protein sequence evolution models that better account for cross-site compositional heterogeneity (and thus fit the data better) were employed (**Fig. S2, S3, S4**). Moreover, our phylogeny supports a derived phylogenetic placement of the photosymbiotic *S. semivirescens*, distant from *P. tenue* and *P. chlorelligera*, suggesting that two spirostomid lineages independently acquired their green algal symbionts (**Fig. 2A**).

**Figure 2.**
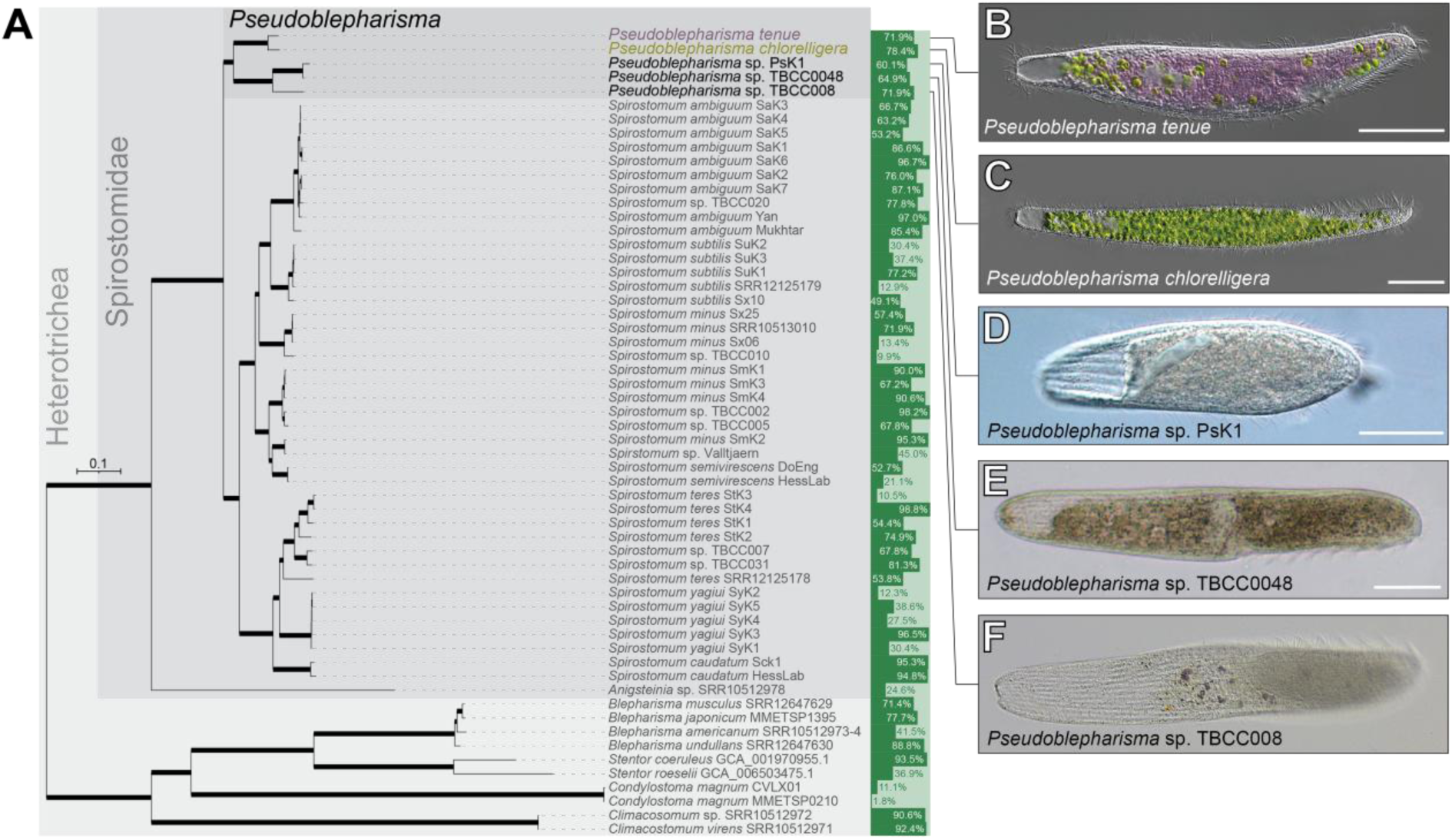
Multiprotein phylogeny of the family Spirostomidae. **A**. Maximum likelihood phylogenetic tree of the Spirostomidae based on 203 proteins with selected heterotrichids as outgroup confidently established the genus *Pseudoblepharisma* as sister to *Spirostomum*. The LG+PMSF+F+G4 model was employed using a guide tree inferred with the LG+C60+F+G4 model. Thickened branches represent branch support values (SH-aLRT, UFBoot2+NNI, and non-parametric bootstrap) higher than 70%. See Fig. S4 for details. Bars represent genome/transcriptome completeness measured with BUSCO scores. **B-F**. Representative light micrographs of *P. tenue* (**B**), *P. chlorelligera* (**C**), *Pseudoblepahrisma* sp. PsK1 (**D**), *Pseudoblepahrisma* sp. TBCC0048 (**E**), and *Pseudoblepharisma* sp. TBCC008 (**F**). Scale bars: 50 µm.

**Figure 3.**
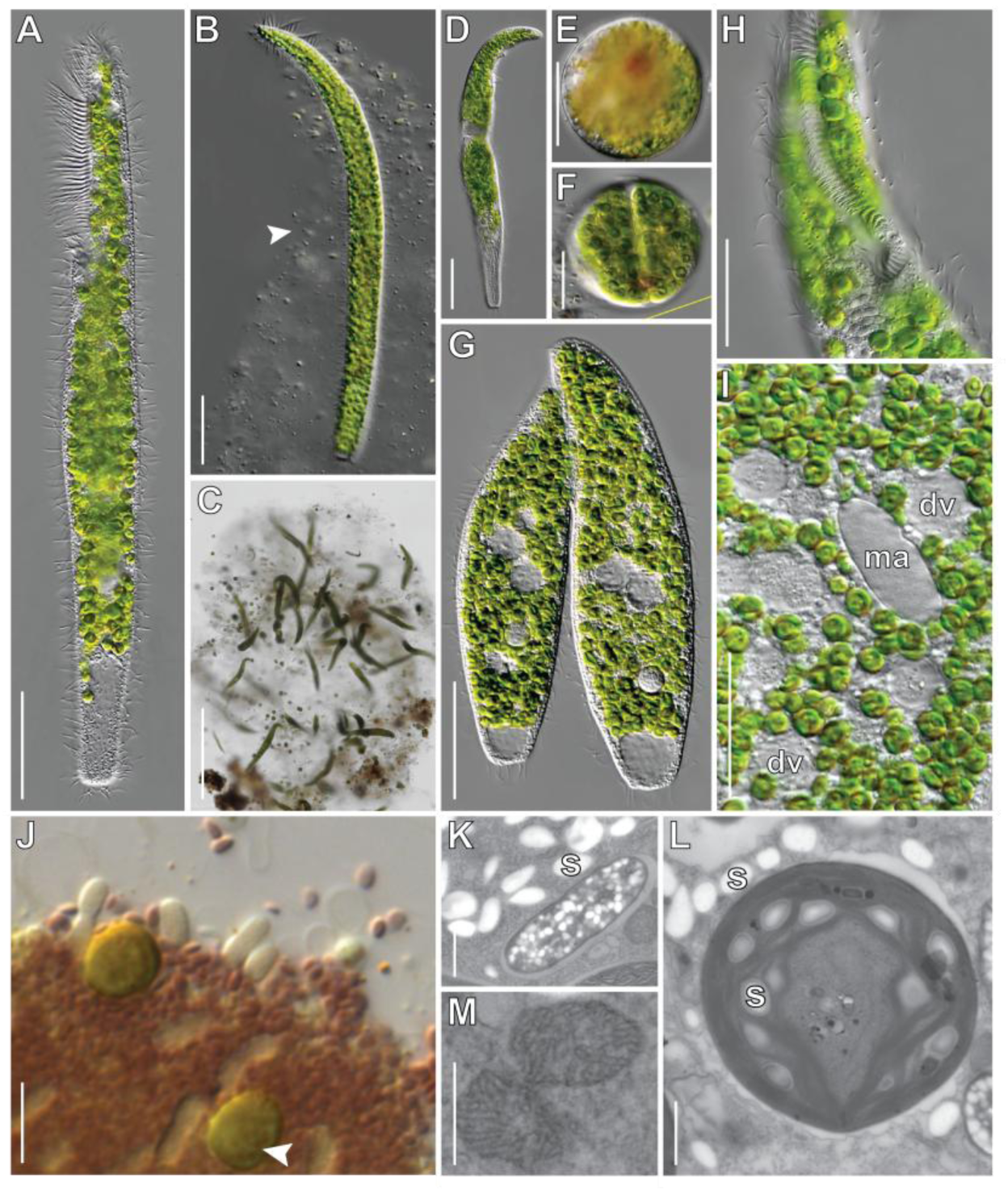
Morphology, life history stages, and ultrastructure of *P. chlorelligera*. **A**. Whole cell. **B**. Mucilaginous lorica (arrowhead) secreted by a whole individual cell. **C**. Macroscopic mucilaginous cluster containing several cells. **D**. Free-swimming cell during division. **E**. Cyst stage. **F**. Cyst undergoing cell division. **G**. Conjugating pair. **H**. Peristome and cytostome. **I**. Detail of cytoplasm with digestive vacuoles (dv) and elliptic macronucleus (ma). **J**. Lugol-stained starch granules in host cytoplasm and green algae (arrowhead), and colorless bacteria. **K**. TEM section of storage granules in the host (starch; s) and colorless bacteria. **L**. TEM section of starch granules (s) in the host cytoplasm and green algal chloroplast. **M**. TEM section of host mitochondria with tubular cristae. Scale bars: 1,000 µm (C), 50 µm (A, B, D, E, G), 25 µm (F, H, I), 5 µm (J), 1 µm (K, M), 0.5 µm (L).

**Figure 4.**
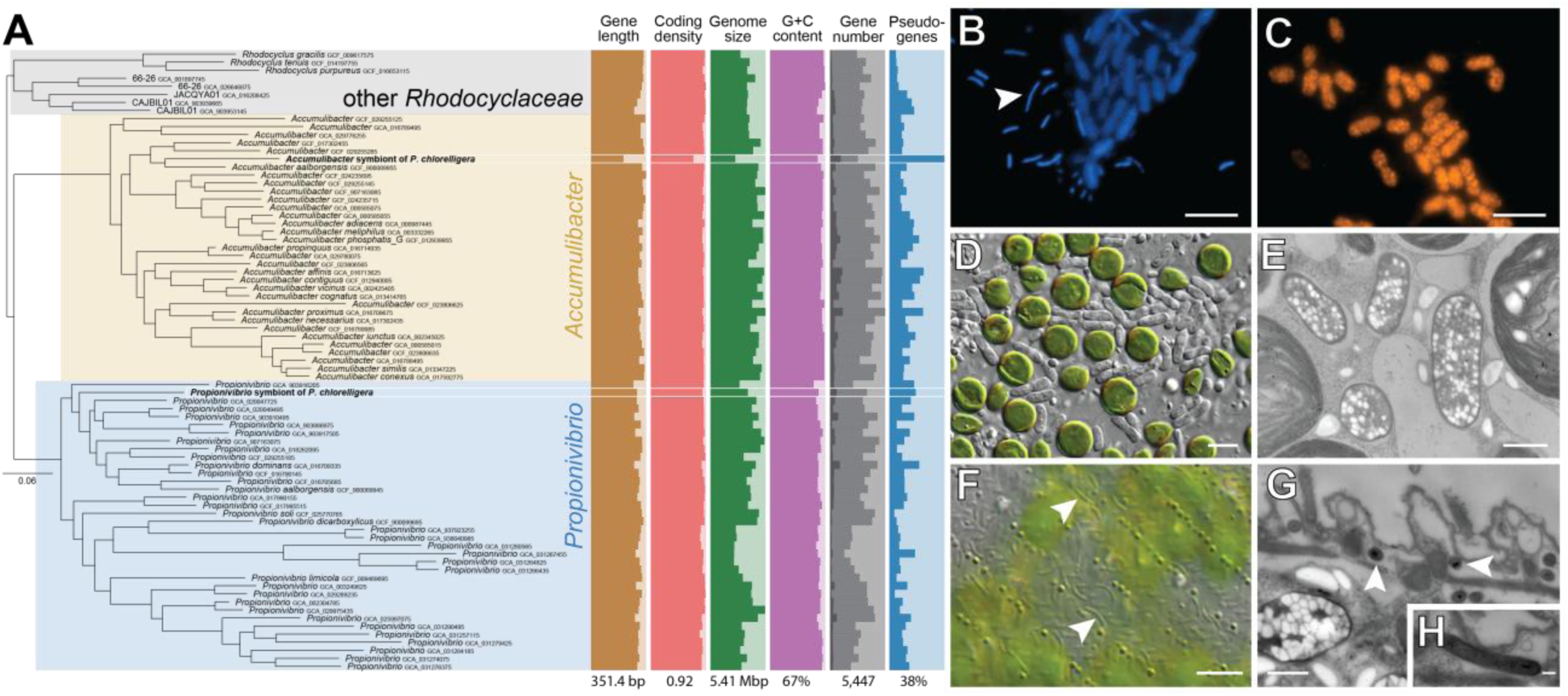
Phylogenetic position, morphology, and cellular localization of bacterial symbionts of *P. chlorelligera*. **A**. Phylogeny of the sister genera *Accumulibacter* and *Propionibrio* within the family *Rhodocyclaceae*. Genome features are displayed next to the phylogenetic tree. Gene length refers to average gene length; genome size is estimated based on completeness measurements with CheckM2 [18]; gene number is decomposed into intact (light gray) and pseudo- (dark gray) genes; pseudogenes were identified with Pseudofinder [19]; number below genome feature distributions indicate maximum values. **B.** Fluorescent micrograph of DAPI-stained “*Ca.* Accumulibacter symbioticus” (thick) and “*Ca.* Propionivibrio subcutaneus” (thin) symbionts. **C.** Fluorescent micrograph of “*Ca.* Accumulibacter symbioticus” from a FISH experiment using a probe specific to the 16S rRNA of the symbiont Accu441-Cy3. **D.** Light micrograph of intracellular green algae and “*Ca.* Accumulibacter symbioticus”. **E.** TEM section of “*Ca.* Accumulibacter symbioticus” with abundant storage granules. Note the symbiosomal membranes surrounding the cells. **F.** Light micrograph of “*Ca.* Propionivibrio subcutaneus” in the cortex of the host cell. **G.** TEM section confirming the cortical localization of “*Ca.* Propionivibrio subcutaneus” symbionts (cross section). **H.** TEM section of a “*Ca.* Propionivibrio subcutaneus” symbiont (longitudinal section). Scale bars: 5 µm (B, C, D, F), 1 µm (E, G), 0.2 µm (H).

### A green-colored *Pseudoblepharisma* species that lacks purple bacterial symbionts

The exact evolutionary relationship between photosymbiotic *Pseudoblepharisma* species, and their endosymbionts, have remained unclear. *P. chlorelligera* (formerly treated as *P. tenue* var. *chlorelligera* [12, 13] or *viride* [10, 14]), for example, was suggested to be an ecological morphotype of *P. tenue* that differentiates in response to varying environmental conditions [14]. A reason for this is that the purely green *P. chlorelligera* and the purple-green *P. tenue* species co-occur in the organic matter-rich sediments of the Simmelried ponds (Allensbach, Germany) (**Fig. 3**). A closer look at the Simmelried population of *P. chlorelligera* reveals clear morphological and ecological differences with respect to *P. tenue*.

The most salient morphological characteristic of *P. chlorelligera* compared to *P. tenue* is the absence of intracellular purple bacteria and the much larger number of green algae. *P. chlorelligera* was packed with intracellular green algae with an average of 352 green algal cells per host (n=6) (**Fig. 3A**). This differs strongly from *P. tenue* that only carries about 19 algal cells per host cell (**Fig. 1A**). *P. chlorelligera* reaches larger cell sizes than *P. tenue* (maximum length ∼360 µm vs. ∼220 µm), and frequently formed mucilaginous loricae (**Fig. 3B**), which sometimes gave rise to macroscopic structures harboring small populations (**Fig. 3C**). Such extracellular mucilaginous housings have never been observed in the purple-green *P. tenue* despite extensive sampling. We also observed dividing cells of *P. chlorelligera*, both in free-swimming (**Fig. 3D**) and cyst (**Fig. 3E, 3F**) stages, as well as conjugating cells in dense populations (**Fig. 3G**).

The cells of *P. chlorelligera* exhibited a conspicuous peristome that occupied about a quarter of the cell’s length and is most likely used for filter-feeding as in other spirostomids (**Fig. 3A, 3H**). Indeed, some individuals contained food vacuoles with partially digested material, suggestive of phagotrophy (**Fig. 3I**). Moreover, the cytoplasm contained densely packed storage granules that stained reddish with iodine solution, which is indicative of amylopectin (**Fig. 3J**). These starch granules are likely polymerized from photosynthate leaked from the green algal symbionts and point to excess carbon in the symbiosis. Transmission electron microscopy confirmed the accumulation of starch in the ciliate cytoplasm and the algal chloroplasts (**Fig. 3K, 3L**), and revealed tubular mitochondrial cristae (**Fig. 3M**) which are characteristic of aerobically respiring ciliates. We conclude that *P. chlorelligera* is a mixotroph that relies on both oxygenic photosynthesis by its green algal symbionts and the consumption of bacteria and small eukaryotes (e.g., *Chlorella*-like cells).

Phylogenetic analyses of the rRNA gene operon confirmed that the Simmelried population of *P. chlorelligera* is genetically very close to the populations found in the subtropical swamps of Florida (**Fig. S1**), suggesting a wide geographic distribution of this taxon. In the Simmelried ponds, the purely green *P. chlorelligera* co-exists with the purple-green *P. tenue*—this was also observed in the Czech-Moravian highlands [12]. Despite co-existing in the same habitats, the morphological (e.g., size and symbiont density), behavioral (e.g., lorica development) and genetic data suggest that *P. tenue* and *P. chlorelligera* occupy distinct ecological niches. There is thus substantial evidence (including genome data, see **Fig. 2A**) that supports the herein proposed existence of two distinct *Pseudoblepharisma* species. Hence, we introduce *Pseudoblepharisma chlorelligera* stat. nov. by elevating the taxonomic rank of *P. tenue* var. *chlorelligera* (see Taxonomy section for details). This species constitutes an important piece of the puzzle to understanding the unique symbiosis of *P. tenue*.

### *Pseudoblepharisma chlorelligera* is a quadripartite symbiosis

High-resolution microscopy revealed that *P. chlorelligera* contains relatively large (∼3.8 µm) colorless bacterial symbionts dispersed throughout its cytoplasm (**Fig. 3H, 3L, 3N**). They differ clearly in size, shape, and color from “*Ca.* Thiodyction intracellulare” (*Chromatiacae*, *Gammaproteobacteria*), which inhabits the cytoplasm of *P. tenue* [3]. Similar colorless bacteria were also reported from the Florida populations [14]. To gain further insights into the identity of these colorless bacterial symbionts and the physiology of the *P. chlorelligera* symbiotic consortium, we sequenced the metagenome and metatranscriptome several individually isolated *P. chlorelligera* cells. This allowed us to recover nearly complete draft genomes for each of the symbiotic partners (**Table S3** and **Fig S6**).

Light microscopy provided clear evidence for one type of bacterial symbiont in *P. chlorelligera* (**Fig. 3H, 3L, 3N**). However, inspection of the metagenome revealed the presence of two nearly complete prokaryotic genome bins (or Metagenome Assembled Genomes; MAGs) with very similar nucleotide compositions but different read coverages in all six sequenced WGA libraries (see **Table S3** and **Methods**). Phylogenetic analyses showed that the bacterial genomes belong to the sister genera *Accumulibacter* and *Propionivibrio* in the family *Rhodocyclaceae* (*Gammaproteobacteria*) (**Fig. 4A**). Whereas the genome of the *Accumulibacter* symbiont is considerably reduced relative to those of its closest relatives, the *Propionivibrio* symbiont shows no or very slight genomic reduction (**Fig. 4A**). This is paralleled in their average gene length, coding density, GC content, total gene number, and fraction of pseudogenes (**Fig. 4A**). This suggests that either the *Accumulibacter* symbiont was acquired first or has experienced a higher rate of genome reduction and erosion compared to the *Propionivibrio* symbiont. This discovery prompted us to search for microscopic evidence for the physical localization of the two bacterial symbionts in *P. chlorelligera*. DAPI (4′,6-diamidino-2-phenylindole) staining confirmed the presence of two morphologically distinct intracellular bacteria, which differed drastically in size (**Fig. 4B**). Interestingly, the two types of bacteria occupied different regions in the host cell, as visualized by transmission electron microscopy. The large bacteria (∼3.5 x 1.3 µm) were scattered throughout the cytoplasm (**Fig. 4D, 4E**), while the thin bacteria (∼2.5 x 0.4 µm) localized to the cell cortex, potentially within the ciliate host’s alveolae (**Fig. 4F, 4G, 4H**). Fluorescence *in situ* hybridization (FISH) with specific probes revealed that the larger bacteria in the cytoplasm correspond to the *Accumulibacter* symbionts (**Fig. 4C**). Thus, *P. chlorelligera* represents a quadripartite symbiosis with green algae and two different colorless bacteria as intracellular symbionts. The latter two are distinct from known genotypes and here described as “*Ca.* Accumulibacter symbioticus” and “*Ca.* Propionivibrio subcutaneus” (see Taxonomic Summary for details).

Species of the genera *Accumulibacter* and *Propionivibrio* are referred to as polyphosphate-accumulating organisms (PAO) and glycogen-accumulating organisms (GAO), respectively [18]. Both genera are often found to co-exist in wastewater environments, which are rich in nutrients and organic matter [18]. Reconstructions of the major metabolic pathways based on the MAGs indicate that both bacterial symbionts of *P. chlorelligera* are chemoheterotrophs that can respire aerobically and ferment. In addition, “*Ca.* Accumulibacter symbioticus” is capable of heterotrophic carbon fixation as it has retained form II rubisco (**Fig. S7**). The genome reduction of “*Ca.* Accumulibacter symbioticus” has led to the loss/pseudogenization of the genes responsible for nitrogen fixation, sulfur cycling, urea utilization, and hydrogenase group 1 and 2, such that only the NiFe-group 3 hydrogenase remains (**Fig. S7**). Furthermore, it has lost succinate dehydrogenase and is, therefore, reliant solely on fumarate reductase, which participates typically in anaerobic respiration, although it can function within aerobic respiration [19] (**Fig. S7**). However, almost all genes involved in amino acid synthesis, fermentation, and complex carbon degradation have been retained (**Fig. S7**). In contrast, “*Ca.* Propionivibrio subcutaneus” has not experienced notable genome reduction, and still possesses the genes required for nitrogen assimilation, sulfur cycling, and urea utilization (**Fig. S7**).

Comparisons of the predicted metabolism of the two bacterial symbionts with that of the *P. chlorelligera* host suggests that amino acid provision to the host could be a potential benefit of the symbiotic exchange, given that the host is auxotrophic for many of the essential amino acids (**Fig. S8**). “*Ca.* Propionivibrio subcutaneus” could additionally provide organic nitrogen as ammonia to both the host and “*Ca.* Accumulibacter symbioticus” (**Fig. S9**). Meanwhile, “*Ca.* Accumulibacter symbioticus” has retained the complete heme biosynthesis pathway, in particular *hemC* and *hemH*, that is incomplete in both the host and “*Ca.* Propionivibrio subcutaneus” and could be indicative of metabolic complementation (**Fig. S10, S11**).

Polyphosphate-accumulating organisms typically take up phosphates in oxic conditions and synthesize poly-β-hydroxyalkanoates, most commonly poly-β-hydroxybutyrate (PHB), using the polyphosphates as reducing equivalents in anoxic conditions [20]. PHB is a carbon storge polymer synthesized from acetyl-CoA, often with acetate as the carbon source [21]. The success of polyphosphate-accumulating organisms in microbial communities arises, therefore, when there are fluctuations between oxic and anoxic conditions [22]. In the latter, PHB synthesis was suggested to fulfill an important additional role as an alternative electron sink because it regenerates NAD(P)+ and so relieves potential TCA cycle inhibition [23, 24]. “*Ca.* Accumulibacter symbioticus” has retained the necessary PHB biosynthesis genes (*phbA-C*) (**Table S5**), and the presence of PHB granules in the cytoplasm of “*Ca.* Accumulibacter symbioticus” was verified experimentally with Nile Blue A staining (**Fig. S12**). The accumulation of PHB indicates that “*Ca.* Accumulibacter symbioticus” were in hypoxic or anoxic conditions, which aligns with other aspects of their metabolism, such as their reliance on fumarate reductase.

### The green algal symbionts were acquired in the ancestor of *P. tenue* and *P. chlorelligera*

We investigated the exact phylogenetic provenance of the green algal symbionts using the nuclear rRNA genes (18S and 5.8S), internal transcribed spacer regions (ITS1 and ITS2) of the rRNA gene operon, and the chloroplast *rbcL* gene. The phylogenetic trees reveal that the green algal symbionts of all photosymbiotic spirostomids (i.e. *P. tenue*, *P. chlorelligera*, and *S. semivirescens*) are identical, and most closely related to *Chlorella* sp. K10 (Chlorellaceae, Chlorophyta), an endosymbiont of the freshwater green cnidarian *Hydra viridissima* [25] (**Fig. 5A, S13**). This is consistent with a less densely sampled chloroplast *rbcL* gene phylogeny (**Fig. 5B. S14**; [3]). As the family Spirostomidae is primarily composed of heterotrophs [26–28], it appears unlikely that the common ancestor of the Spirostomidae harbored intracellular algae as symbionts. According to our multiprotein phylogeny, this scenario would require five independent losses of a photosymbiotic lifestyle (**Fig. 2A and Fig. S1**). Given the large phylogenetic distance between photosymbiotic spirostomids, and the prevalence of lateral transfers of *Chlorella*-like symbionts among hosts [29], it is most probable that *Chlorella* sp. K10 was acquired twice by two independent lineages, namely *S. semivirescens* and the ancestor of *P. tenue* and *P. chlorelligera*. As previously reported, *Chlorella* sp. K10 has a versatile energy metabolism (like other green algae [30–32]), namely the capacity to perform several different types of fermentations alongside oxygenic photosynthesis and aerobic respiration [3]. However, because *P. chlorelligera* appears to prefer the upper layers of the muddy sediments found in the Simmelried pond [13] and accumulates abundant starch granules that are most likely polymerized from photosynthate, the intracellular green algae in *P. chlorelligera* are inferred to actively photosynthesize. This contrasts with *P. tenue* that harbors only a few green algal symbionts which, we hypothesize, may primarily ferment in the more anaerobic niche of their symbiotic consortium (see below).

**Figure 5.**
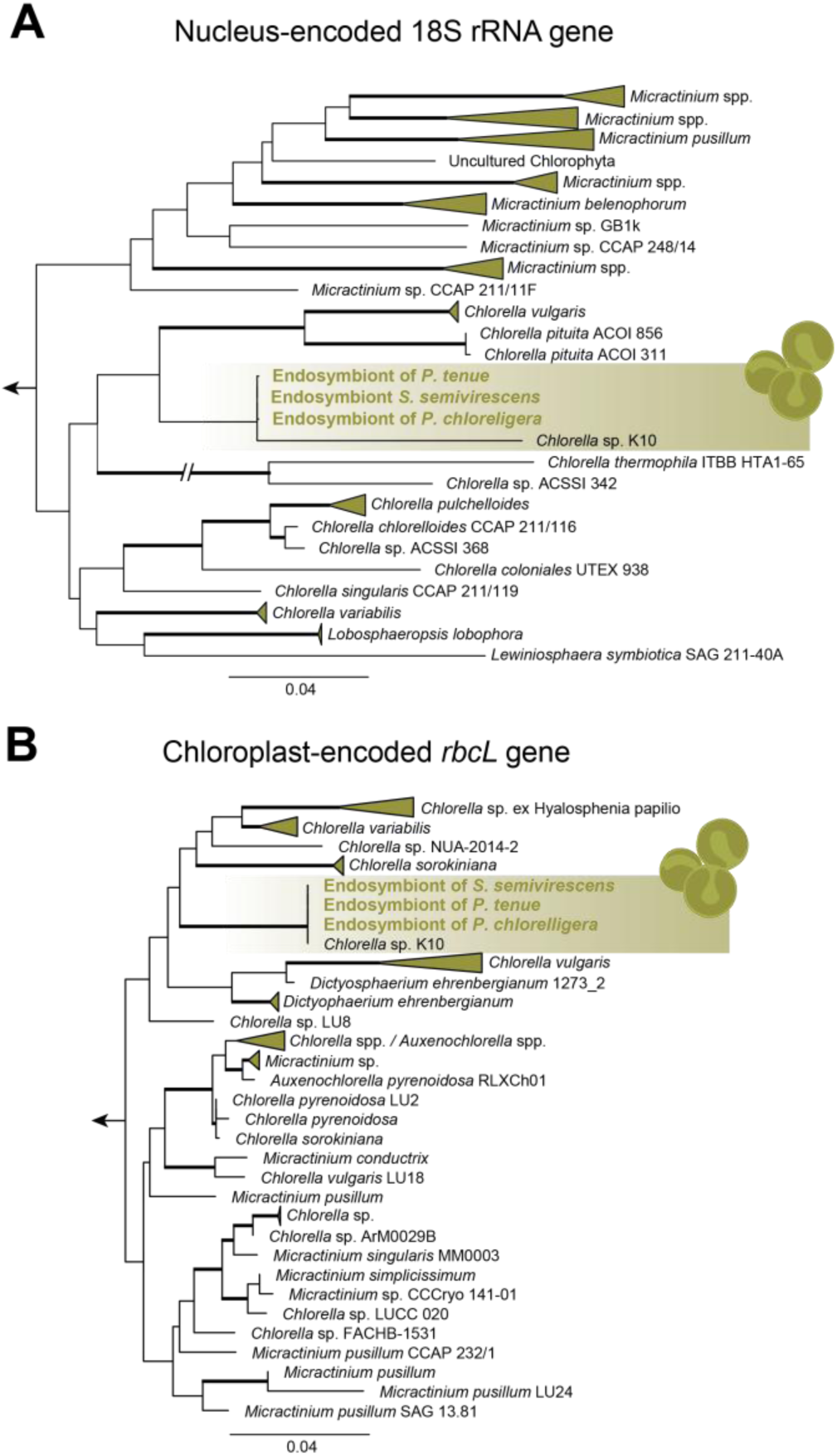
Close phylogenetic association between green algal symbionts in the Spirostomidae and the *Hydra*-symbiont *Chlorella* sp. K10. A. Phylogenetic tree of the Chlorellaceae based on the nucleus-encoded 18S rRNA gene and 132 taxa. B. Phylogenetic tree of the Chlorellaceae based on the chloroplast-encoded *rbcL* gene and 199 taxa. Trees differ in phylogenetic coverage and representation due to differential marker sampling on NCBI GenBank. Only a subtree containing the clade of interest is shown. For complete phylogenetic trees, see Fig. S13 and S14. Support values are SH-aLRT, UFBoot2+NNI, and non-parametric bootstrap. Thickened branches represent support value combinations of greater than or equal to 70%/70%/70%.

### The hypothetical origin of the unique purple-green symbiosis of *Pseudoblepharisma tenue*

Photosymbioses with purple bacteria such as that of *P. tenue* are extremely rare. This poses the question of what factors prevent and favor the establishment of such photosymbioses in nature. It may simply be that a series of improbable events were required for the evolutionary assembly of purple photosymbioses. For example, the pre-symbiotic partners may be confined to a few environments or niches that only partially overlap in nature (e.g., those that provide the right electron donors for anoxygenic photosynthesis). This restricts the opportunities for the partners to co-exist in the same micro-habitat and thus the probability for the evolutionary emergence of the symbiosis. Indeed, purple photosymbioses appear to only thrive in a few environments that provide a favorable, yet still unidentified, set of ecological factors [1]. Based on our new findings, we can now propose an evolutionary narrative for how the unique symbiosis of *P. tenue* was evolutionarily assembled.

We hypothesize that the purely heterotrophic and aposymbiotic ancestor of *P. tenue* and *P. chlorelligera* inhabited freshwater sediments that were depleted or low in oxygen. In support of this, both *Pseudoblepharisma* and *Spirostomum* species are well known to dwell in oxygen-poor sediments (e.g., [33]). *P. chlorelligera*, for example, was found at dissolved oxygen amounts as low as 0.24% saturation [14]. Such hypoxic microhabitats offer an ecological opportunity to facultative anaerobes that is not available to faster-growing and more speciose aerobes, e.g., lower competition for prey bacteria [34]. In low-oxygen environments, *Pseudoblepharisma* and *Spirostomum* species likely rely on fumarate respiration for energy conservation. This energy metabolic pathway consists of a short electron transport chain where electrons extracted from NADH by complex I (NADH dehydrogenase) are transferred to reduce fumarate into succinate via rhodoquinol and a reversed complex II (succinate dehydrogenase). The protons pumped into the intermembrane space by complex I contribute to the proton motive force which is then used by complex V (an F_1_F_O_-ATP synthase) to synthesize ATP [35]. We previously showed that *P. tenue* has the potential to perform fumarate respiration [3]. An extended analysis of the distribution of anaerobic energy metabolism reveals that fumarate respiration is ubiquitous not only among *Pseudoblepharisma* and *Spirostomum* species [16], but also among other heterotrichids making it likely ancestral to the entire Heterotrichea (**Fig. S15**). This contrasts with anaerobic ciliates in the Intramacronucleata which are more specialized to an anaerobic lifestyle and instead rely on an oxygen-sensitive Fe-Fe hydrogenase for ATP production in their hydrogenosomes (**Fig. S15**) [36, 37]. Altogether, these analyses suggest that the *Pseudoblepharisma* ancestor was a facultative anaerobe that may have resembled in physiology and lifestyle the non-photosymbiotic colorless *Pseudoblepharisma* species discovered here (i.e., TBCC008, TBCC0048 and PsK1; **Fig. 2B, S1, S2, S3, S4**). This lifestyle is likely a prerequisite for the origin of purple photosymbioses as purple sulfur bacteria (such as the ancestor and free-living relatives of “*Ca*. Thiodictyon intracellulare”) are anaerobic photosynthesizers with a preference for anoxic habitats [38], and often rely on hydrogen sulfide which is poisonous to aerobic eukaryotes by inhibiting cytochrome *c* oxidase [39].

*P. tenue* constitutes a dual photosymbiosis as it harbors two contrasting photosynthetic symbionts. As such, one symbiont may have taken residence inside its host before the other. The phylogenetic evidence suggests that the green algae were acquired in the common ancestor of *P. tenue* and *P. chlorelligera*. The green algal symbionts thus preceded the acquisition of purple bacteria in *P. tenue* and colorless bacteria in *P. chlorelligera*. These green algal symbionts were potentially acquired from *Chlorella* sp. K10 cells that had escaped or been released from another eukaryote host and were thus pre-adapted to a symbiotic lifestyle. The initial selective advantage of the green algal symbionts may have been to provide carbon from oxygenic photosynthesis, as in a classical photosymbiosis. However, if the *Pseudoblepharisma*—*Chlorella* symbiosis had a preference for hypoxic sediments and/or was exposed to prolonged darkness, its physiology may have turned anaerobic where both ciliate host and green algal symbionts increasingly relied on fermentation for ATP production—green algae switch to fermentation and hydrogen production under anoxic conditions both in the light and the dark [30–32] (**Fig. 6**). This created a conducive metabolic environment for the establishment of a second photosymbiotic relationship by making electron donors other than sulfide (e.g., acetate, propionate, or molecular hydrogen; products of host and green algal fermentation), as well as a nitrogen source (e.g., amino acids or ammonia), readily available to the pre-symbiotic purple bacteria that entered the host cell via phagocytosis (**Fig. 6**). This intracellular environment, in turn, allowed for the subsequent loss of sulfur dissimilation and nitrogen fixation in “*Ca*. Thiodictyon intracellulare” (**Fig. 6**). Although equipped with the same photosynthetic capacity at first (oxygenic photosynthesis by *Chlorella*), the ancestors of *P. tenue* and *P. chlorelligera* followed divergent evolutionary paths and formed very different photosymbioses that can still co-exist in the same habitat (**Fig. 6**).

**Figure 6.**
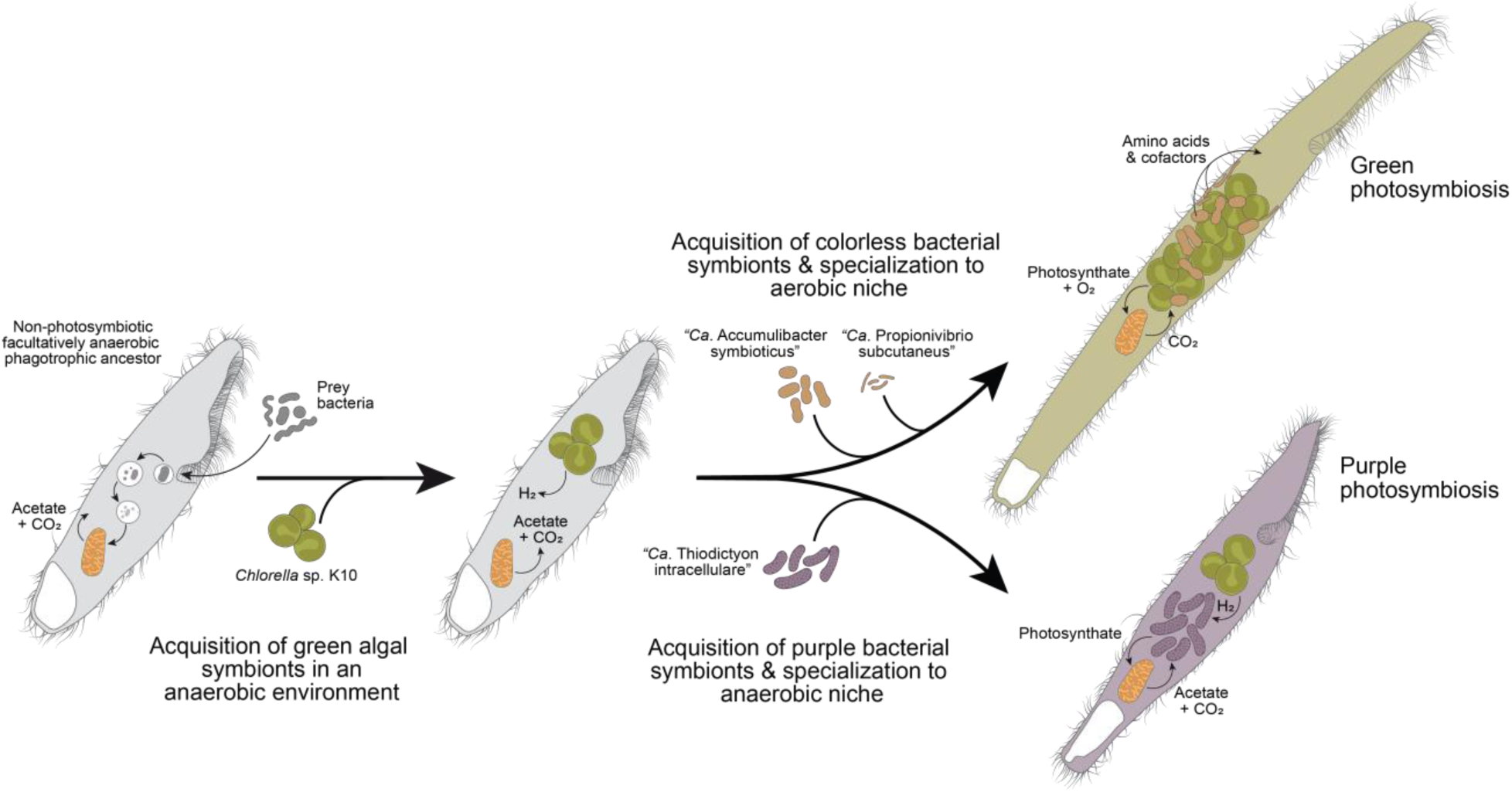
Hypothetical sequence of events that led to the origin of the purple-green photosymbiosis of *P. tenue*. See main text for details. Note that all ancestors and extant species remain phagotrophic (or mixotrophic in the case of photosymbiosis).

This study has revealed the order of endosymbiont acquisition in *P. tenue* and potential pre-adaptations that facilitated the symbiosis. These insights have, in turn, shed light on the question of how two contrasting photosynthetic symbionts came to exist within the same host. Moreover, we have expanded the known diversity of the genus *Pseudoblepharisma* by discovering new non-photosymbiotic species and confidently establishing the sister relationship between the genera *Pseudoblepharisma* and *Spirostomum* in the family Spirostomidae. Our genome-based predictions on the physiology of photosymbiotic *Pseudoblepharisma* species represent the basis for future experimental work on the identity of the metabolites that are exchanged among symbiotic partners.

### Taxonomy

**Phylum Ciliophora** DOFLEIN, 1901

**Class Heterotrichea** STEIN, 1859

**Order Heterotrichida** STEIN, 1859

**Family Spirostomidae** STEIN, 1867

**Genus *Pseudoblepharisma*** KAHL, 1926

***Pseudoblepharisma chlorelligera*** stat. nov.

#### Authorship

MUÑOZ-GÓMEZ, KREUTZ & HESS

#### Basionym

*Pseudoblepharisma tenue* var. *chlorelligera* ŠRÁMEK-HUŠEK, 1949

#### Revised description

Slender, weakly contractile cells, 200-360 µm long, ∼30 µm wide, with pointed anterior end and truncated posterior. Peristome about ¼ of the cell body length, with adoral membranelles but without undulating membrane. About 20 longitudinal kineties. Contractile vacuole posterior with inconspicuous dorsal collecting channel, flattened after systole. Macronucleus localized in the mid to posterior region of the cell, ellipsoidal, with several small, adherent/embedded micronuclei. Cells densely packed with ∼350 symbiotic algae (*Chlorella* sp.) of about 6-7 µm and colonized by two types of heterotrophic bacteria. Cells can swim at 50-100 µm s^-1^ under anti-clockwise rotation (as seen from anterior end) and, when undisturbed, form hyaline, gelatinous loricae/tubes (semi-sessile lifestyle). Cysts spherical with smooth wall, about 60 µm in diameter, potentially associated with cell division.

#### Differential diagnosis

Differs from *P. tenue* in larger cell length, hundreds of endosymbiotic *Chlorella* cells, absence of intracellular purple bacteria, and presence of gammaproteobacterial endosymbionts. Differs from *Spirostomum semivirescens* in smaller cell length and macronuclear morphology (ellipsoidal vs. moniliform).

Because the newly identified gammaproteobacterial endosymbionts could not be grown in pure culture, the requirements of the ICNP [40] for valid publication—including deposition of a viable type strain—cannot be fulfilled. Following the guidelines of Murray and Stackebrandt [41] for uncultured prokaryotes, we provide a provisional taxonomic description and assign the *Candidatus* status to these symbiotic bacteria:

**Phylum *Pseudomonadota*** OREN et al., 2021

**Class *Gammaproteobacteria*** STACKEBRANDT et al., 1988

**Order *Rhodocyclales*** GARRITY et al., 2006

**Family *Rhodocyclaceae*** GARRITY et al., 2006

**Genus “*Candidatus* Accumulibacter”** HESSELMANN et al., 1999

**“*Candidatus* Accumulibacter symbioticus”** [(Rhodocyclales, Gammaproteobacteria) NC; NA; R; NAS (GenBank Number), G (MAG Accession); S (*Pseudoblepharisma chlorelligera*, Ciliophora; cytoplasm); M; probably aer. resp. or ferment. (reduced N-fixation, S-metabolism, urea utilization, H2-ases, and SDH genes lost, but Rubisco form II and heterotrophic potential retained)]. **Authorship:** MUÑOZ-GÓMEZ, KREUTZ & HESS, (REF). **Etymology:** *symbioticus* (N.L. masc. adj.) = symbiotic; referring to the obligate intracellular lifestyle within *Pseudoblepharisma chlorelligera*. **Additional information:** Large (∼3.5×1.3 μm), colorless, intracellular bacteria scattered throughout the host cytoplasm. Each cell bounded by at least two membranes (possibly three), one of which may be a vacuolar membrane, packed with electron-translucent storage granules discernible in TEM micrographs; no other life history stages observed. Genome strongly reduced relative to free-living *Accumulibacter* spp.; loss of genes for nitrogen fixation, sulfur metabolism, urea utilization, hydrogenases (groups 1–2) and succinate dehydrogenase (SDH); retention of Rubisco form II and broad heterotrophic metabolic potential; aerobic respiration and fermentation predicted. Genetically defined by the 16S rRNA gene sequence XXXXXXXXXX and the draft genome XXXXXXXXXX (both GenBank). Reference material: Resin block for TEM (XXXXXXXXXX; deposited in the “Protists Collection” at the Department of Life Sciences of the National History Museum in London (Cromwell Road, UK) containing a single host cell collected in pond II of the “Simmelried” moorland, Constance, Germany; 47.717767, 9.09375.

**Genus *Propionivibrio*** TANAKA et al., 1991**“*Candidatus* Propionivibrio subcutaneus”** [(Rhodocyclales, Gammaproteobacteria) NC; NA; R; NAS (GenBank Number), G (MAG Accession); S (*Pseudoblepharisma chlorelligera*, Ciliophora; intracellular/potentially alveolae); M; potentially chemoheterotrophic (N-assimilation, S-cycling, urea utilization, and complete respiratory systems retained); aer. resp. or ferment.]. **Authorship:** MUÑOZ-GÓMEZ, KREUTZ & HESS, (REF). **Etymology:** *subcutaneus* (N.L. masc. adj.) = “lying beneath the surface”, referring to the cortical localization in the host cell. **Additional information:** Slender (∼2.5×0.4 μm), colorless, intracellular bacteria located in the cortex of the host cell, potentially within the alveolae. Each cell bounded by two membranes (in addition to alveolar membrane), devoid of discernible storage granules; no other life history stages observed. The genome is not markedly reduced; it retains pathways for nitrogen assimilation, sulfur cycling, urea utilization, and complete respiratory systems. Chemoheterotrophic metabolism with capability for aerobic respiration and fermentation predicted. Genetically defined by the 16S rRNA gene sequence XXXXXXXXXX and the draft genome XXXXXXXXXX (both GenBank). Reference material: Resin block for TEM (XXXXXXXXXX; deposited in the “Protists Collection” at the Department of Life Sciences of the National History Museum in London (Cromwell Road, UK) containing a single host cell collected in pond II of the “Simmelried” moorland, Constance, Germany; 47.717767, 9.09375.

## Methods

### Sampling and cultivation

*Pseudoblepharisma chlorelligera* was sampled in the Simmelried region as follows. Water and sediment samples were taken from pond II (naming according to [10]) of the Simmelried peat bog area near Hegne, Germany (47.717767, 9.09375) at a depth of about 30 cm. These samples were transported in 200-mL jars and lastly combined into 1-L bottles with a loose screw cap to allow the formation of a bottom sediment and different zones. These bottles were kept at 15°C under dim, indirect light with a 14:10-hour light/dark cycle. Heterotrophic strains of *Spirostomum* and *Pseudoblepharisma* were sampled at diverse locations as follows. Water samples of several hundred mL were taken from the organic sediment of the ponds and transferred to the laboratory. From these samples several individuals (>20) of the same ciliate morphotype were picked and transferred to Erlenmeyer flasks with 75 mL Volvic mineral water supplemented with 1 mL lettuce extract, 1 mL soil extract [42] and three halved grains of naked barley. These non-monoclonal cultures were grown at 15 °C in the dark.

### Transmitted light microscopy

For high-resolution imaging of live cells, two microscopes were used: an Olympus BX 50 microscope and a Zeiss IM35 inverted microscope, both equipped with differential interference contrast optics, high-resolution objective lenses, and an electronic flash. Digital images were taken with an Olympus E-P5 digital camera and Canon EOS 6D digital single lens reflex camera, respectively, and processed with Adobe Photoshop CS4 (Adobe Systems).

### Transmission electron microscopy

About 30 cells of *P. chlorelligera* were isolated from natural samples and collected in about 200 µl of distilled water. This cell suspension was quickly injected into 200 µl of freshly prepared fixative [80 µl of 25% glutaraldehyde, 200 µl of 4% osmium tetroxide solution, and 120 µl of half-strength medium Waris-H (pH 7) with 1.5 mM Hepes], resulting in a final concentration of 2.5% glutaraldehyde and 1% osmium tetroxide. Cells were incubated for 1 min at room temperature and for another 45 min at 6°C, then quickly washed with distilled water (twice, supernatant was removed after cells settled), and dehydrated by a series of acetone:water mixtures (30, 50, 80, and 90%) and pure acetone (twice) for 5 min each at −20°C. Acetone was then replaced by 50% EPON in acetone, incubated for 24 hours at about 6°C for infiltration, and then the tube was opened and stored for another 24 hours in a fume hood to let the acetone evaporate. The sample was then mixed with fresh EPON and cured at 65°C for 1.5 days. Single ciliate cells were sectioned (∼60 nm thickness) with a Leica EM UC7 ultramicrotome (Leica Microsystems), stained with uranyl acetate (1%, 15 min) and Reynolds’ lead citrate solution (3.5 min), and imaged with a CM10 TEM (FEI Europe Main Office, Eindhoven, The Netherlands) and a Gatan ORIUS SC200W TEM charge-coupled device (CCD) camera (Gatan Inc., Pleasanton, CA).

### Fluorescent *In Situ* Hybridization

Using a 16S rRNA gene sequence obtained by Sanger sequencing that showed >98% identity with *Accumulibacter* species (*Rhodocyclaceae*, *Gammaproteobacteria*), we retrieved potential FISH probes with the “Design Probes” web tool of the software toolset DECIPHER under default hybridization conditions [43]. One of the probe sequences with maximum specificity (score of 0), reasonable hybridization efficiency, and no reported cross-hybridizations with nontarget sequences was analyzed with the online tool “mathFISH” [44] and manually modified to optimize binding to the target sequence. The resulting probe Accu441 (5′-CCC AAG CAA TTT CTT TCC CGC-3′) was lastly checked for specificity in silico with the “probeMatch” database [45] and the SILVA probe match and evaluation tool “TestProbe 3.0.” The probe showed no hits with zero mismatches in both databases and only few hits with one mismatch indicating high specificity (as of March 2020). The Cy3-conjugated probe Accu441-Cy3 was ordered (biomers.net GmbH, Ulm, Germany), and its binding to bacterial cells from *P. chlorelligera* was evaluated under varying formamide concentrations (0 to 60%) using a modification of previously published FISH protocols [46, 47]. In brief, cells of *P. chlorelligera* were isolated from natural samples, washed twice in distilled water, placed on gelatin-coated glass slides (coated with 0.1% gelatin in water), and air-dried. Some ciliate cells were deliberately burst with a pipette tip to spread the endosymbionts. Cells on dry slides were fixed with formaldehyde (3.7% in water) for 10 min, then rinsed in distilled water, and air-dried. To minimize potential autofluorescence, photosynthetic pigments of bacteria and algae were first extracted from the samples by incubating the slides in a methanol:water mixture (8:2) at 65°C for 10 min, then rinsed in ethanol (100%), and air-dried. The cells were then covered with 50 to 100 µl of hybridization buffer containing probe Accu441-Cy3 at an appropriate concentration (5 ng/liter) and 0 to 60% formamide (according to [47]) and then incubated in a moist chamber at 46°C for 90 min. After incubation, the slides were quickly rinsed with warm washing buffer (48°C) and subsequently incubated in washing buffer at 48°C for 25 min. Last, slides were rinsed with distilled water, air-dried, and mounted with SlowFade Diamond Antifade Mountant (Thermo Fisher Scientific).

Routine epifluorescence microscopy of FISH samples was done with the Zeiss Axiophot microscope equipped with phase contrast optics, the Zeiss filter set 43 HE (excitation, 550/25 nm; emission, 605/70 nm), and the camera AxioCam HRc (Carl Zeiss). For confocal laser scanning microscopy, a Leica TCS SPE system and the Leica Confocal Software (Leica Microsystems) were used. The samples were excited with lasers (561 nm for Cy3 and 488 nm for chlorophyll) and scanned in three dimensions with step sizes of < 1 µm. Confocal laser scanning microscopy (CLSM) stacks were processed and analyzed with the software LAS AF lite (Leica Microsystems) and the image processing package Fiji [48]. This included the selection of pseudocolors for fluorescence channels, the adjustment of brightness, and the combination of different channels (merged channels) and/or of several focal planes (Z-projections).

### Staining

Staining with Nile Blue A was done as follows. Individually picked *P. chlorelligera* cells were burst with a custom-made glass Pasteur pipette and heat-fixed on glass slides. Enough Nile Blue A 1% solution was added to cover the cells, and the slide was then incubated at 55°C for 10 min [49]. This was followed by rinsing with ultra-pure water and incubating with 8% acetic acid for 1 min. DNA staining was done by adding DAPI 30 µM to heat-fixed cells on a slide and incubating at 4°C for 1 h. For Lugol staining, several ciliate cells were isolated with a glass micropipette and transferred to a wet-mount preparation. A small droplet of 2% Lugol’s iodine was placed at the edge of the cover slip, allowing the solution to be drawn into the mount. The cells were then gently compressed by removing excess liquid and examined using bright-field and differential interference contrast microscopy.

### Single-cell PCR

Individual cells from cultured isolates or environmental populations of *Spirostomum*-like ciliates (**Table S1**) were isolated using glass micropipettes, washed twice in sterile distilled water, and transferred into a maximum of 10 µl of nuclease-free water in a 0.2-mL tube. These tubes were then snap-frozen in liquid nitrogen and stored at −20°C. To extract genomic DNA (gDNA) from isolated cells, the following components were added to the frozen tubes in order: 50 µl of tris-HCl buffer (10 mM; pH 8.5), 50 µl of Chelex 100 (5%; Sigma-Aldrich), and 2 µl of proteinase K (20 mg/ml; Thermo Fisher Scientific). The solution was then mixed by vortexing for 15 s, briefly centrifuged, and then incubated at 56°C for 45 min and lastly at 98°C for 20 min. After incubation, the samples were centrifuged at 17,000g for 3 min and stored at 4°C. PCR of single-cell gDNA extract was performed using Q5 DNA Polymerase (New England Biolabs) following the manufacturer’s protocol. A volume of 4 µl of gDNA extract was used per 50-µl PCR reaction mix. To amplify most of the rRNA gene operon (comprising the 18S rRNA gene, ITS1-5.8S rRNA-ITS2, and the D1D2 region of the 28S rRNA gene) of the ciliate host, the primers Euk_A (5′-AAC CTG GTT GAT CCT GCC AG-3′) and D1D2rev2 (5′-ACG ATC GAT TTG CAC GTC AG-3′) were used [50, 51]. The PCR cycle was as follows: 98°C for 30 s (1x), 98°C for 10 s, 66°C for 15 s, 72°C for 70 s (35x), and 72°C for 120 s (1x). PCR amplicons were purified with the NucleoSpin Gel and PCR Clean-up Kit (Takara Bio) and sequenced with the Sanger sequencing method (Eurofins Genomics, Ebersberg, Germany) using the primers EukA (5′-AAC CTG GTT GAT CCT GCC AG-3′) and EukB (5′-TGA TCC TTC TGC AGG TTC ACC TAC -3′). Sanger reads were assembled on Benchling (https://benchling.com). Overlapping consensus sequences of the ciliate rRNA operon were assembled in Geneious www.geneious.com). **Table S1** lists the accession numbers of the Sanger-sequenced 18S rRNA genes.

### Single-Cell Whole Genome and Transcriptome Amplification and sequencing

Single cells from environmental samples or isolate cultures were isolated using a modified Pasteur glass micropipette, washed, and deposited in 0.2-mL tubes as detailed above were inspected under low magnification to confirm cell integrity and then snap-frozen in liquid nitrogen. The REPLI-g Advanced DNA Single Cell Kit (QIAGEN) was used to perform a single-cell whole-genome amplification (WGA) on three individual ciliates. A 65°C incubation step for 10 min was introduced to facilitate lysis of endosymbionts. The lid was set to a temperature of 70°C. The REPLI-g WTA Single Cell Kit (QIAGEN) was used to perform single-cell whole transcriptome amplification. A combination of random and oligo dT primers were used for the symbiotic consortium of *P. chlorelligera*, whereas only oligo dT primers were used for all other isolates (see **Table S2**). Illumina sequencing libraries were made with the Illumina TruSeq kit from the products of all three WGA variations. The premium whole genome amplification protocol (SQK-LSK109) was followed for WGAs from cells of *P. chlorelligera* for Nanopore long-read sequencing. Illumina short-read (2x150 bp) and Oxford Nanopore long-read sequencing were done on a NovaSeq and PromethION instruments at the Cologne Center for Genomics at the University of Cologne, Germany. **Table S2** lists the accession number for each of the libraries generated and sequenced as part of this study.

### Metagenomics of *P. chlorelligera*

Raw Illumina short reads were quality-trimmed and -filtered with AfterQC v. 0.9.6 (options: -f -1 - t -1) [52]. The processed reads were subsequently normalized by down-sampling with BBNorm v.39.01 (options: mindepth=5 target=50) prior to assembly. The raw Nanopore long reads were processed with Porechop v. 0.2.4 to remove adapters (options: --discard_middle), quality-trimmed and -filtered with NanoFilt v. 2.8.0 (options: --headcrop 50 -q 8 -l 1000), and quality-filtered once more using reference Illumina reads with Filtlong v. 0.2.1 (options: --keep_percent 90 --trim --split 500 --length_weight 10 --min_length 1000). The length and quality of the processed Nanopore long reads were assessed with NanoPlot v. 1.41.6. The normalized Illumina short reads from four libraries were co-assembled with Nanopore long reads from two libraries with metaSPAdes v3.15.5 (options: --sc) [53].

Assembled scaffolds were processed through the metagenomic pipeline in Anvi’o v7 [54]. In brief, scaffolds shorter than 1,000 bp were discarded, and reads from each library were separately mapped on to the scaffolds with Bowtie 2 v. 2.5.1 [55] to obtain their coverages. Taxonomic affiliations were assigned to each scaffold with Kaiju v.1.9.2 [56] and the pre-built index for the nr_euk database. Scaffolds were clustered according to their composition and coverage and manually inspected with anvi-interactive [54]. Genome bins for the ciliate host and green algae were manually identified using scaffold taxonomy assignments, and nucleotide composition and/or mean coverage. Gene annotation was performed with BlastKOALA and KofamKOALA [57].

### Binning of *Accumulibacter* and *Propionivibrio* MAGs

A composite bin of both *Accumulibacter* and *Propionivibrio* was refined manually with anvi-refine [54] by removing foreign scaffolds. The refined composite bin was further binned with MyCC [58] using coverage profile from six different metagenomic libraries and the 4mer setting (see **Table S3**). To generate coverage profiles, both Illumina short and Nanopore long reads were mapped to the composite bin scaffolds with Bowtie 2 [55] and Minimap2 [59], respectively. SAMtools [60] was then used to sort and index the resulting bam files. MyCC binning yielded two well separated clusters that corresponded to nearly complete *Accumulibacter* and *Propionivibrio* genomes with little redundancy. Completeness and redundancy were estimated with CheckM2 [61]. Taxonomic affiliation of each binned was assigned with GTDB-Tk v1.5.0 [62]. Gene prediction and annotation was done with Bakta [63], BlastKOALA [57], KofamKOALA [57], and METABOLIC [64]. The prediction of pseudogenes was done with Pseudofinder v1.1.0 [65]. The quantification of prokaryotic transcripts was done with FADU [66]. All gene annotations were combined into a single table for interpretation.

### Metabolic pathway prediction and reconstruction for *Accumulibacter* and *Propionivibrio* MAGs

For the gene retention and metabolic pathway analysis, the predicted pseudogenes were first removed from the datasets. The gene retention/loss comparison across the *Rhodocyclaceae* was based on the ‘FunctionHit’ METABOLIC output, which calculates the presence/absence of functional sets of proteins associated with core metabolic functions (**Table S5**). The absence of critical genes was verified by targeted BLASTp searches (*Accumulibacter*: *mdh*, *CS* and *hisN*. *Propionivibrio*: *CbiJ, hisN, AcnB, fumA, ATPF0* and *ATPF1* subunits. *P. chlorelligera*: *coq7, ATPeF0* and *ATPeF1* subunits). The detailed amino acid and vitamin comparisons were made from the ‘KEGGModuleStepHit’ METABOLIC output that calculates the presence/absence of each step within a KEGG module. The vitamin analysis includes the results for all modules in the ‘Cofactor and vitamin metabolism’ category, and the amino acid analysis all the modules in the categories: ’Arginine and proline metabolism’, ’Cysteine and methionine metabolism’, ’Branched-chain amino acid metabolism’, ’Histidine metabolism’, ’Lysine metabolism’, ’Serine and threonine metabolism’, ’Aromatic amino acid metabolism’ and ’Other amino acid metabolism’. The KEGG pathway maps were made with KEGG mapper color tool (v5) [67], using the annotated ‘KEGG_ID’ column (**Table S4**). RStudio (v2025.09.0+387) running R (v4.5.1) was used for additional analysis and figure generation [68, 69].

### Phylogenetic analyses

The 18S rRNA sequences of the new *Spirostomum*/*Pseudoblepharisma* isolates were added to the rRNA gene operon (18S rRNA-ITS1-5.8S rRNA-ITS2-D1D2 28S rRNA) dataset used in our previous study ([3]; itself based on [28]). 18S rRNA genes from publicly available transcriptomes and some WGAs produced in this study were reconstructed using phyloFlash v3.4.2 [70]. Sequences were aligned with MAFFT v7.525 G-INS-i (--globalpair --maxiterate 1000) and trimmed with TrimAl v1.4.rev15 (-automated1). Phylogenetic trees were reconstructed with IQ-TREE v2.2.6 (-allnni -pers 0.3 -nstop 500; [71]) using the MFP+MERGE of ModelFinder (-m MFP-MERGE -rcluster 10; [72]) to find the best-fit partitioning scheme (18S:TNe+I+R2; ITS1:GTR+F+I+G4; 5.8S:TNe+I+R2; ITS2:GTR+F+I+G4; D1D2-28S:GTR+F+I+G4; COI:GTR+F+I+G4).

The 18S and 5.8S rRNA genes, and ITS1 and ITS2 regions, of the *Chlorella*-like symbionts of *P. tenue*, *P. chlorelligera*, and *S. semivirescens* were identified through BLASTn searches [73]using the *Meyerella* sp. strain ACSSI 362 (OL619998.1) as a query against their draft genome assemblies. The scaffolds containing these markers were subsequently searched against the Rfam database to reliably identify the boundaries of the 18S and 5.8S rRNA genes. The rRNA gene operon was manually annotated in Geneious (www.geneious.com). These sequences, as well as the 18S rRNA genes of *Chlorella* symbionts of Hydra reported previously [25], were then added to the 18S-ITS1-5.8S-ITS2 alignment of Krivina et al. (2022) [74]. Phylogenetic trees were reconstructed with IQ-TREE v2.2.6 (-allnni -pers 0.3 -nstop 500; [71]) using the MFP+MERGE of ModelFinder (-m MFP-MERGE -rcluster 10; [72]) to find the best-fit partitioning scheme (18S: TNe+R3; ITS1:GTR+F+I+G4; 5.8S: TNe+R3; ITS2:GTR+F+I+G4).

Chloroplast *rbcL* gene homologs in nucleotide sequence were identified through tBLASTx searches [73]. Sequences with >50% query coverage that were taxonomically assigned to “Chlorellaceae” or “unassigned Chlorellaceae” were retained for further analysis. Aligned tBLASTx regions were extracted and aligned as nucleotide sequences based on their protein alignments (i.e., codon-based) with MACSE [75] using genetic code 11 (The Bacterial Archaeal and Plant Plastid Code). Non-homologous ends were removed manually in AliView [76]. Phylogenetic trees were reconstructed with IQ-TREE v2.2.6 (-allnni -pers 0.3 -nstop 500; [71]) using the MFP+MERGE of ModelFinder (-m MFP-MERGE -rcluster 10; [72]) to find the best-fit partitioning scheme (GTR+F+I+R4).

Genome surveys and transcriptomes publicly available were downloaded and assembled for all species that belong to the Heterotrichea in NCBI GenBank as of March 2023. Raw Illumina short reads from WTA libraries and public RNA-Seq experiments were quality-trimmed and -filtered as above, and subsequently assembled with Trinity v2.15.1 [77]. Completeness assessment was done with BUSCO v5.2.2 [78]. Proteome prediction was done with TransDecoder v5.7.1 (-m 50). WGA libraries were assembled with SPAdes (options: --sc) [53]. *de novo* repeat identification was done with RepeatModeler v2.0.3 and subsequent annotation and masking using RepeatMasker v4.1.5. Proteome prediction from genome assemblies (derived from WGA Illumina libraries; see **Table S2**) was done with BRAKER v 2.1.6 pipeline [79, 80]. The retrieval and curation of orthologs, and dataset construction, was done by following the PhyloFisher pipeline [81]. In short, the PhyloFisher pipeline performs similarity searches to obtain candidate homologs and add them to a curated database of eukaryotic taxa (limited to the Ciliophora here) which are then used to build single-gene phylogenetic trees that are inspected and annotated manually to label orthologs, paralogs, and contaminants. The curated gene orthologs were then process with PREQUAL [82] to remove non-orthologous regions, aligned with MAFFT [83], trimmed with Divvier [84] and trimAl [85], and finally concatenated into a supermatrix. Phylogenetic inference was done with IQ-TREE v2.2.6 using the LG4X and LG+C60+G4+F models. The inferred LG+C60+G4+F tree was subsequently used as a guide for phylogenetic inference with the LG+PMSF+G4+F model [86]. Support values were merged with the -sup option in IQ-TREE [71]. Visualization of phylogenetic trees with FigTree v1.4.4 and iTOL v6 [87].

Close relative genomes of the *Accumulibacter* and *Propionivibrio* symbionts were identified from the GTDB database r220 using GTDB-Tk v2.1.1 [62]. These genomes were retrieved and a phylogenomic dataset was built with GToTree v1.8.6 and the Gammaproteobacteria marker protein set [88]. Phylogenetic inference was subsequently done with IQ-TREE v2.3.5 [71] and Q.pfam+C20+F+R4 model.

### Similarity searches of metabolic energy enzyme among ciliates

Putative homologs of anaerobic energy-metabolism enzymes were identified in ciliate proteomes using reciprocal best-hit (RBH) similarity searches. For each reference species, query proteins retrieved from previous studies [35, 36, 89] were searched against individual ciliate proteomes using DIAMOND BLASTp v2.1.16 [90]. For each query, the top-scoring hit in the target proteome (lowest E-value, highest bit score) was retained. These candidate hits were then searched back against the full proteome of the original reference species. Protein pairs were considered reciprocal best hits if each sequence recovered the other as its top-scoring match in the forward and reverse searches. Only reciprocal pairs passing additional quality filters (pairwise amino-acid identity ≥25%, query and subject alignment coverage ≥70%, and protein length ratio between 0.75 and 1.25) were retained as putative homologs. All searches used an E-value cutoff of 1×10^−5^.

## Acknowledgements

This work was supported by a NASA Exobiology Program grant (80NSSC24K1875) to SAM-G and the Emmy Noether Programme of the German Research Foundation (grant 417585753) to SH. M.E.S.S. received support from a College of Life Science fellowship from the Wissenschaftskolleg zu Berlin. We thank Ragib Ahsan (University of Ulsan) for helping with the identification of ciliate species isolated from South Korea, and Michael Müller (Regenstauf, Germany) and Michael Butkay (Hannover, Germany) for natural samples and picked ciliate cells. NGS experiments were conducted at the Cologne Center for Genomics.

## Data availability

Raw sequencing reads have been deposited at NCBI SRA under the BioProject PRJNA1000070. See **Table S2** for SRA accessions.

## Supplementary figures legends

**Figure S1. Phylogeny of the Spirostomidae based on the rRNA gene operon and mitochondrial COI gene.** The tree is rooted between the genus *Anigsteinia* and all other taxa. See Methods for details. Support values are SH-aLRT (%) / ultrafast bootstrap (%) / non-parametric bootstrap (%). Only branch support values higher than 70% / 70% / 70% are displayed.

**Figure S2. Phylogeny of the Spirostomidae based on 203 proteins and the LG4X model.** The tree is rooted on a selection of non-spirostomid heterotrichids. Support values are SH-aLRT (%) / ultrafast bootstrap (%). Only branch support values higher than 70% / 70% are displayed.

**Figure S3. Phylogeny of the Spirostomidae based on 203 proteins and the LG+C60+G4+F model.** The tree is rooted on a selection of non-spirostomid heterotrichids. Support values are SH-aLRT (%) / ultrafast bootstrap (%). Only branch support values higher than 70% / 70% are displayed.

**Figure S4. Phylogeny of the Spirostomidae based on 203 proteins and the LG+PMSF(C60)+G4+F model.** The tree is rooted on a selection of non-spirostomid heterotrichids. Support values are SH-aLRT (%) / ultrafast bootstrap (%). Only branch support values higher than 70% / 70% are displayed.

**Figure S5. Micrograph plate of *Pseudoblepharisma* sp. PsK1 (Ulsan, South Korea). A.** Whole cell body view of specimen 1. **B.** Whole cell body view of specimen 2. **C.** Closeup of peristome, and adoral membrane (am), and cytostome. **D.** Close up view of ciliary rows and cortical granules. **E.** Close up view of ciliary rows. **F.** Oval macronucleus (ma). **G.** Posterior contractile vacuole (cv) and digestive vacuoles (arrowheads). Scale bars: 55 µm (A, B), 10 µm (C, D, E, F, G).

**Figure S6. Summary of the metagenome of *P. chlorelligera*.** Anvi‘o diagram of the metagenome of the *P. chlorelligera* symbiotic consortium. Scaffolds are organized according to their tetranucleotide composition. The coverage values correspond to four short-read Illumina libraries (WGA3, WGA4, WGA5, and WGA6) made from three different *P. chlorelligera* single cells. The colored taxonomy ticks correspond to contigs whose genes are labeled “Ciliophora”, “Chlorophyta”, and “Candidatus Accumulibacter” and “Propionivibrio” based on similarity searches.

**Figure S7. Comparison of the presence/absence of core metabolic functions between the bacterial endosymbionts *P. chlorelligera* and their relatives.** In the grid, each column is a core metabolic function with the gene names provided as the column name and each row is a species in the *Rhodocyclaceae*. The dark purple shaded boxes depict the presence of a function in the genome, while white/light purple depict absence. The *Accumulibacter* and *Propionivibrio* symbionts of *P. chlorelligera* are highlighted in light purple. The functions are grouped into metabolic categories that are highlighted in different colors at the top of the columns. The phylogenetic relationships of the species are shown to the right of the grid.

**Figure S8. Comparison of the key steps in amino acid metabolism across *P. chlorelligera* and its bacterial endosymbionts.** The amino acid metabolic pathways are listed across the x-axis and each KEGG metabolic step per pathway on the y-axis.

**Figure S9. KEGG map of nitrogen metabolism across *P. chlorelligera* and its bacterial endosymbionts.** The gene product boxes are colored purple if present in the *Propionivibrio* symbiont, green if present in the *Accumulibacter* symbiont, and orange if present in *P. chlorelligera* host.

**Figure S10. Comparison of the key steps in cofactor and vitamin synthesis across *P. chlorelligera* and its bacterial endosymbionts.** The cofactor and vitamin metabolic pathways are listed across the x-axis and each KEGG metabolic step per pathway on the y-axis.

**Figure S11. KEGG map of porphyrin metabolism across *P. chlorelligera* and its bacterial endosymbionts.** The gene product boxes are colored purple if present in the *Propionivibrio* symbiont, green if present in the *Accumulibacter* symbiont, and orange if present in *P. chlorelligera* host.

**Figure S12. Evidence of PHB granules in the *Accumulibacter* symbiont.** Nile Blue A staining *Accumulibacter* symbionts released from the cytoplasm of *P. chlorelligera*. Arrowheads point to fluorescently labeled PHB granules. See Methods for details. Scale bar: 5 µm.

**Figure S13. Phylogeny of the Chlorellaceae based on the 18S and 5.8S rRNA genes and ITS1 and ITS2 regions.** The tree is rooted in between *Parachlorella* clade and all other taxa. See Methods for details. Support values are SH-aLRT (%) / ultrafast bootstrap (%) / non-parametric bootstrap (%). Only branch support values higher than 70% / 70% are displayed.

**Figure S14. Phylogeny of the Chlorellaceae based on the chloroplast *rbcL* gene.** The tree is rooted in its midpoint. See Methods for details. Support values are SH-aLRT (%) / ultrafast bootstrap (%) / non-parametric bootstrap (%). Only branch support values higher than 70% / 70% are displayed.

**Figure S15. Phylogenetic distribution of anaerobic energy metabolic enzymes in representative species of the Ciliophora.** Candidate homologs were identified based on reciprocal best-hit (RBH) similarity searches. Dark grey corresponds to detected RBHs and white to absence. Light grey corresponds to conserved complexes that are inferred to be present but had no RBH most likely due to incomplete transcriptome/genome data. Red corresponds to large phylogenetic blocks of absent RBHs.

**Table S1.** *Spirostomum-* and *Pseudoblepharisma*-like cells sampled in this study.

| Strain | Location | Accession number (18S rRNA gene) |
| --- | --- | --- |
| *Pseudoblepharisma chlorelligera* | Germany | XXXXXXXXXX |
| *Pseudoblepharisma* sp. PsK1 | South Korea | XXXXXXXXXX |
| *Pseudoblepharisma* sp. TBCC008 | Germany | XXXXXXXXXX |
| *Pseudoblepharisma* sp. TBCC048 | Germany | XXXXXXXXXX |
| *Spirostomum semivirescens* HessLab | Germany | XXXXXXXXXX |
| *Spirostomum* sp. TBCC0010 | Germany | XXXXXXXXXX |
| *Spirostomum* sp. TBCC0016 | Germany | XXXXXXXXXX |
| *Spirostomum* sp. TBCC0017 | Germany | XXXXXXXXXX |
| *Spirostomum* sp. TBCC0018 | Germany | XXXXXXXXXX |
| *Spirostomum* sp. TBCC002 | Germany | XXXXXXXXXX |
| *Spirostomum* sp. TBCC0020 | Germany | XXXXXXXXXX |
| *Spirostomum* sp. TBCC0022 | Germany | XXXXXXXXXX |
| *Spirostomum* sp. TBCC0024 | Germany | XXXXXXXXXX |
| *Spirostomum* sp. TBCC0025 | Germany | XXXXXXXXXX |
| *Spirostomum* sp. TBCC0027 | Germany | XXXXXXXXXX |
| *Spirostomum* sp. TBCC0030 | Germany | XXXXXXXXXX |
| *Spirostomum* sp. TBCC0031 | Germany | XXXXXXXXXX |
| *Spirostomum* sp. TBCC0032 | Germany | XXXXXXXXXX |
| *Spirostomum* sp. TBCC0033 | Germany | XXXXXXXXXX |
| *Spirostomum* sp. TBCC0034 | Germany | XXXXXXXXXX |
| *Spirostomum* sp. TBCC0035 | Germany | XXXXXXXXXX |
| *Spirostomum* sp. TBCC0036 | Germany | XXXXXXXXXX |
| *Spirostomum* sp. TBCC0037 | Germany | XXXXXXXXXX |
| *Spirostomum* sp. TBCC0038 | Germany | XXXXXXXXXX |
| *Spirostomum* sp. TBCC0040 | Germany | XXXXXXXXXX |
| *Spirostomum* sp. TBCC0041 | Germany | XXXXXXXXXX |
| *Spirostomum* sp. TBCC005 | Germany | XXXXXXXXXX |
| *Spirostomum* sp. TBCC006 | Germany | XXXXXXXXXX |
| *Spirostomum* sp. TBCC007 | Germany | XXXXXXXXXX |

**Table S2.** Single-cell genome and transcriptome data generated as part of this study. *Species that do not have an associated 18S rRNA gene in Table S1.

| Species name | Location | Type of library | Accession number | Sequencing technology | Size (GB) |
| --- | --- | --- | --- | --- | --- |
| *Pseudoblepharisma chlorelligera* | Simmelried, Germany | WGA | SRR25464810 | Illumina | 30.22 |
| *Pseudoblepharisma chlorelligera* | Simmelried, Germany | WGA | SRR25464809 | Illumina | 89.20 |
| *Pseudoblepharisma chlorelligera* | Simmelried, Germany | WGA | SRR25464798 | Illumina | 12.63 |
| *Pseudoblepharisma chlorelligera* | Simmelried, Germany | WGA | SRR25464794 | Illumina | 73.80 |
| *Pseudoblepharisma chlorelligera* | Simmelried, Germany | WGA | SRR25464793 | Nanopore | 7.10 |
| *Pseudoblepharisma chlorelligera* | Simmelried, Germany | WGA | SRR25464792 | Nanopore | 14.52 |
| *Pseudoblepharisma chlorelligera* | Simmelried, Germany | WTA | SRR25464791 | Illumina | 63.32 |
| *Pseudoblepharisma chlorelligera* | Simmelried, Germany | WTA | SRR25464790 | Illumina | 19.22 |
| *Pseudoblepharisma chlorelligera* | Simmelried, Germany | WTA | SRR25464789 | Illumina | 34.05 |
| *Pseudoblepharisma* sp. PsK1 | Ulsan, South Korea | WTA | SRR25464806 | Illumina | 10.79 |
| *Pseudoblepharisma* sp.<br>TBCC008 | Rheinbach,<br>Germany | WTA | SRR25464808 | Illumina | 33.01 |
| *Pseudoblepharisma* sp.<br>TBCC008 | Rheinbach,<br>Germany | WTA | SRR25464788 | Illumina | 50.11 |
| *Pseudoblepharisma* sp.<br>TBCC048 | Regenstauf,<br>Germany | WGA | SRR25464807 | Illumina | 32.69 |
| *Spirostomum*<br>*semivirescens* HessLab | Hemmingen,<br>Germany | WGA | SRR25464797 | Illumina | 28.84 |
| *Spirostomum caudatum*<br>HessLab* | Simmelried,<br>Germany | WTA | SRR25464799 | Illumina | 28.82 |
| *Spirostomum teres* StK2* | Samcheok,<br>South Korea | WTA | SRR25464796 | Illumina | 18.78 |
| *Spirostomum teres* StK4* | Ulsan, South<br>Korea | WTA | SRR25464795 | Illumina | 51.82 |
| *Spirostomum* sp.<br>TBCC002 | Kassel,<br>Germany | WTA | SRR25464805 | Illumina | 27.70 |
| *Spirostomum* sp.<br>TBCC005 | Heimerzheim,<br>Germany | WTA | SRR25464804 | Illumina | 57.52 |
| *Spirostomum teres*<br>TBCC007 | Cologne<br>(Hürth),<br>Germany | WTA | SRR25464803 | Illumina | 75.12 |
| *Spirostomum* sp.<br>TBCC010 | Brühl,<br>Germany | WTA | SRR25464802 | Illumina | 78.87 |
| *Spirostomum ambiguum*<br>TBCC020 | Heimerzheim,<br>Germany | WTA | SRR25464801 | Illumina | 86.87 |
| *Spirostomum* sp.<br>TBCC031 | Hannover,<br>Germany | WTA | SRR25464800 | Illumina | 42.92 |

**Table S3.** Mean coverage values for each symbiont genome bin across six different sequenced WGA libraries.

|  | **Completeness**<br>**(BUSCO/CheckM2)** | **WGA3** | **WGA4** | **WGA5** | **WGA6** | **WGA7** | **WGA8** |
| --- | --- | --- | --- | --- | --- | --- | --- |
| ***Pseudoblepharisma chlorelligera*** | 87.9%<br>(alveolata_odb12) | 305.45 | 2.34 | 122.83 | 388.32 | 76.18 | 155.82 |
| ***Chlorella* sp. K10** | 81.4%<br>(chlorophyta_odb12) | 8.73 | 104.65 | 2.43 | 6.69 | 3.12 | 4.54 |
| ***“Ca. Accumulibacter symbioticus”*** | 95.2%<br>(CheckM2) | 16.60 | 206.04 | 8.58 | 32.80 | 25.08 | 38.00 |
| ***“Ca. Propionivibrio subcutaneus”*** | 98.04%<br>(CheckM2) | 6.97 | 57.58 | 2.36 | 6.68 | 3.65 | 5.25 |

Table S4. Functional annotations of the predicted proteins of *P. chlorelligera* and its bacterial endosymbionts.

Table S5. Predicted KEGG metabolic functions of *Accumulibacter* and *Propionivibrio.* symbionts and their relatives. This table is the result of a METABOLIC analysis which is itself the basis for Fig. S7, S8 and S10.

## Notes

### Competing Interest Statement

The authors have declared no competing interest.

