## Supplementary FIgures for "The phylogenetic context for the origin of a unique purple-green photosymbiosis"

### Supplementary Figure 1

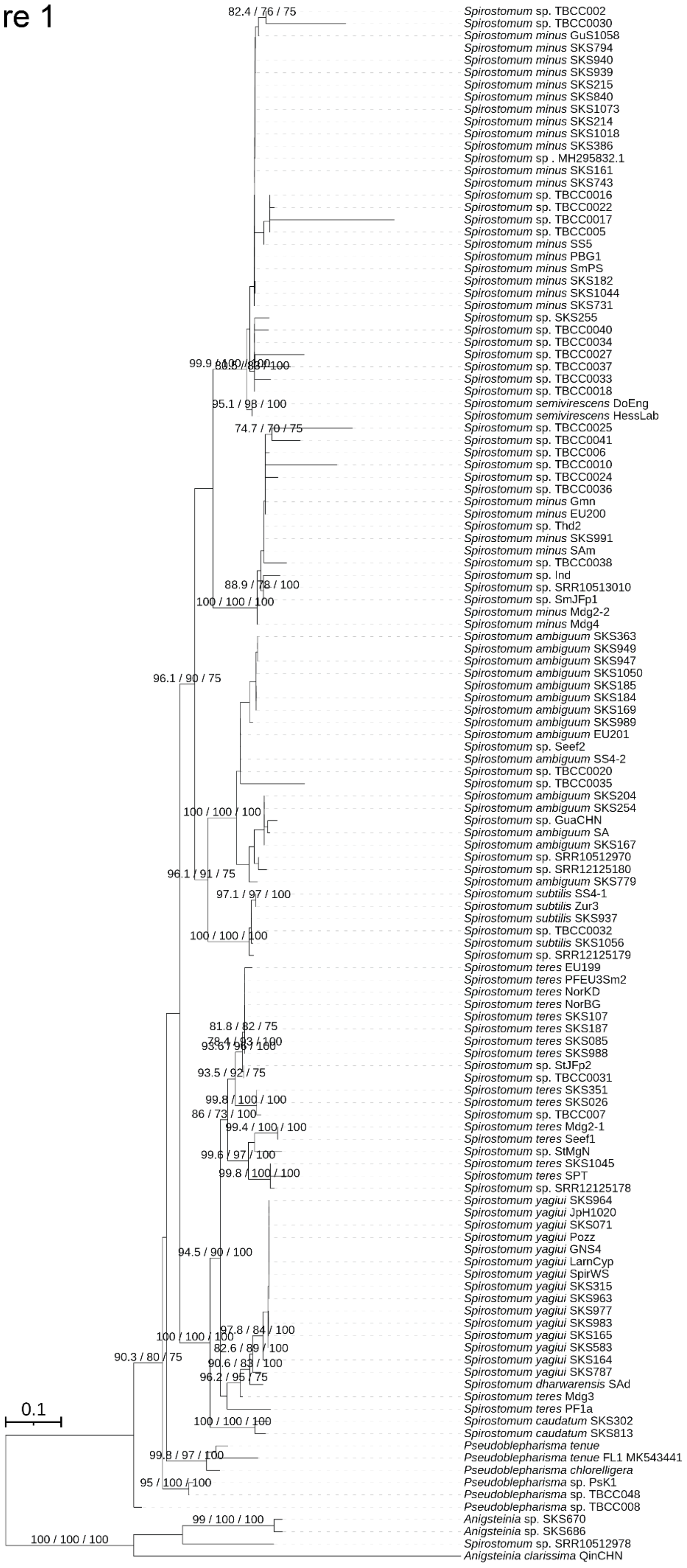

### Supplementary Figure 2

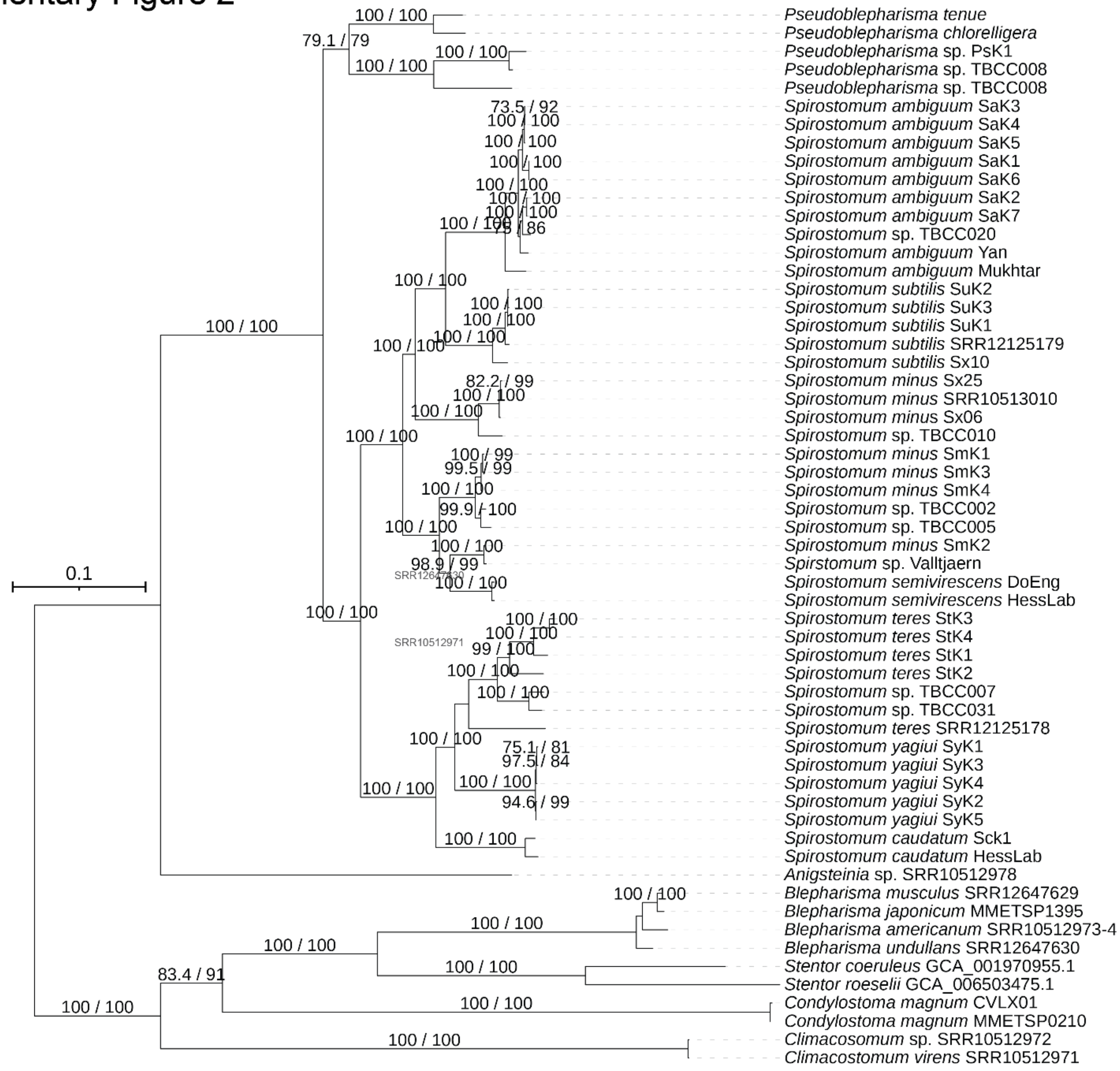

#### Supplementary Figure 3

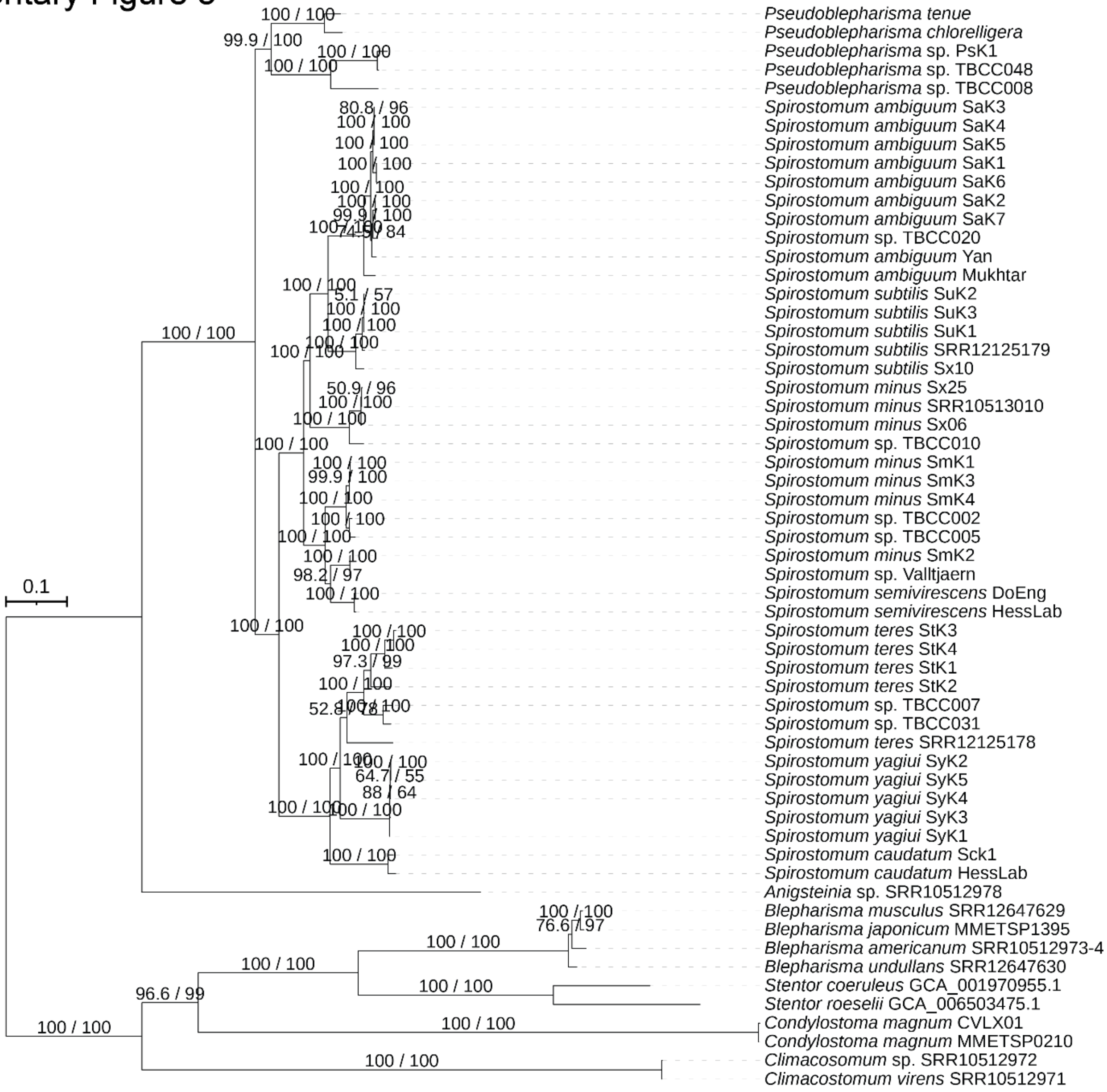

Supplementary Figure 4

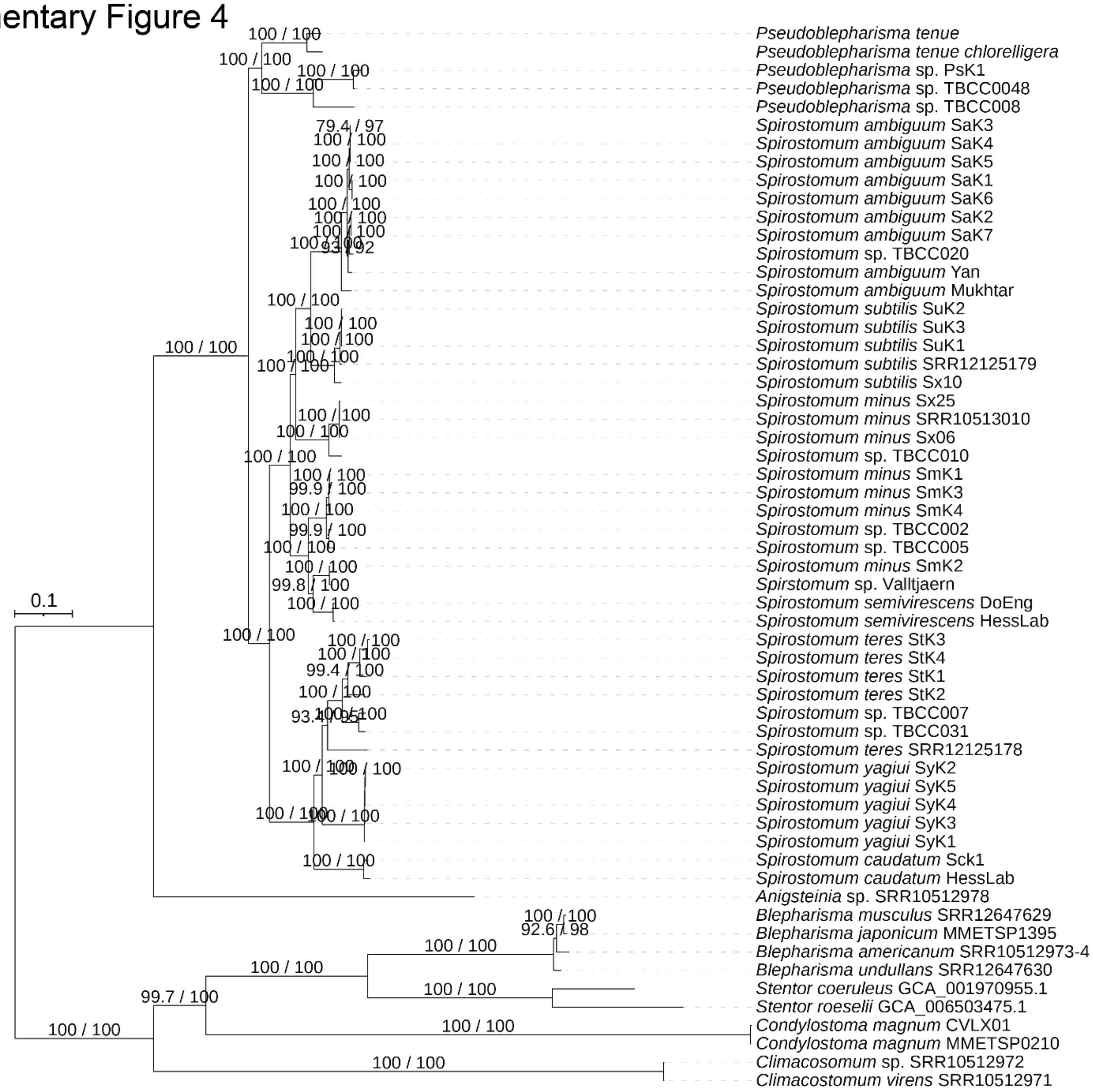

Supplementary Figure 5

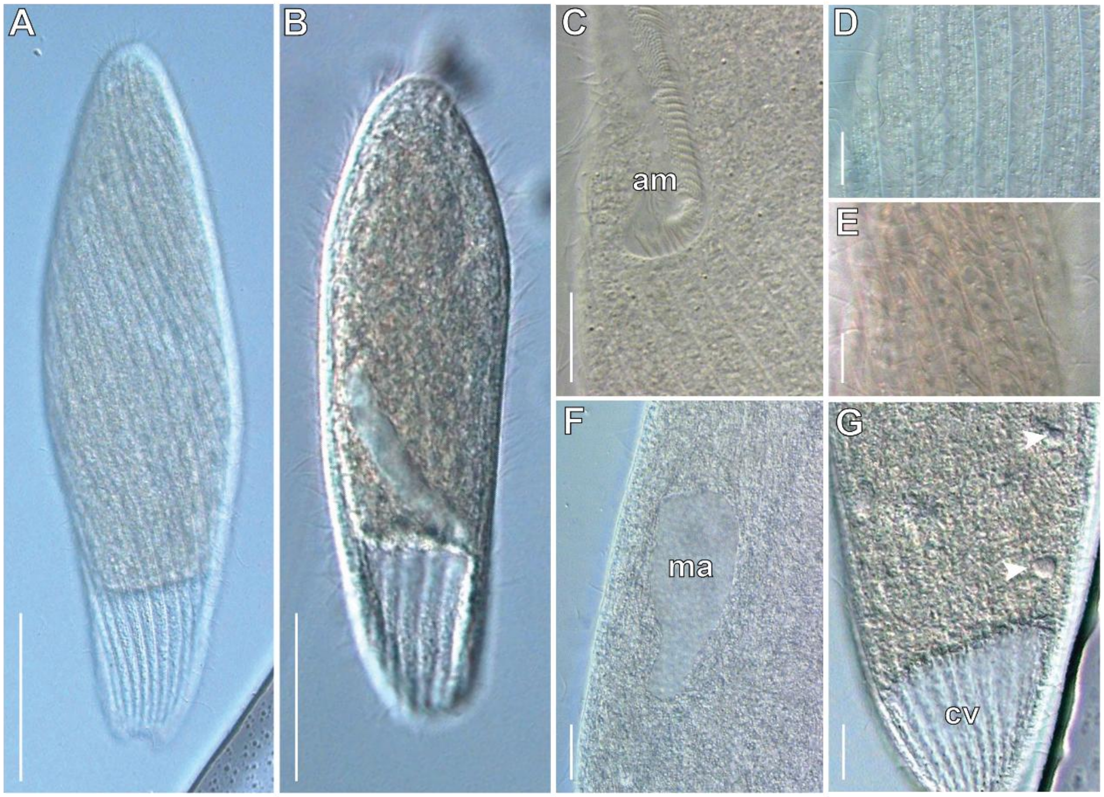

#### Supplementary Figure 6

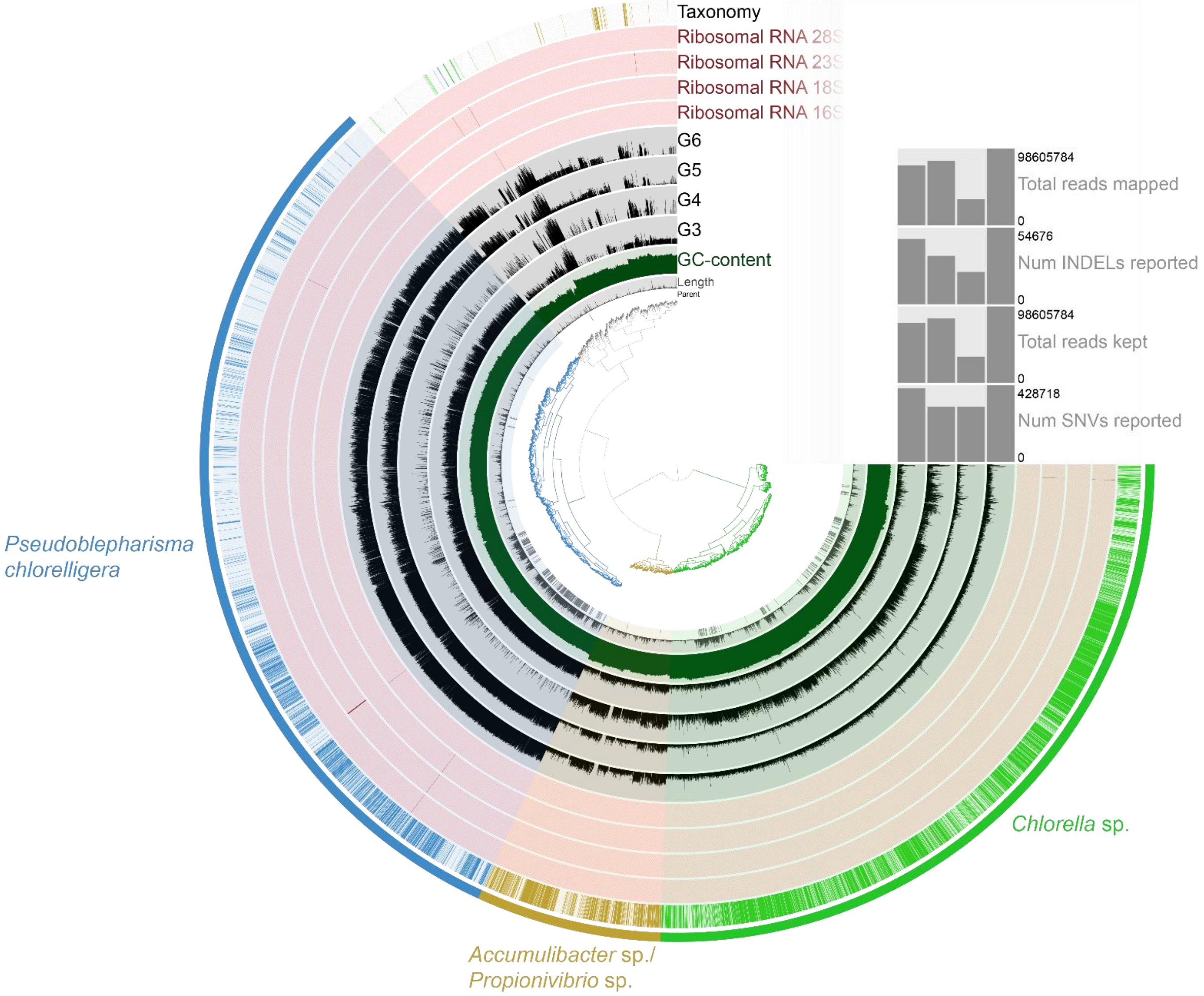

### Supplementary Figure 7

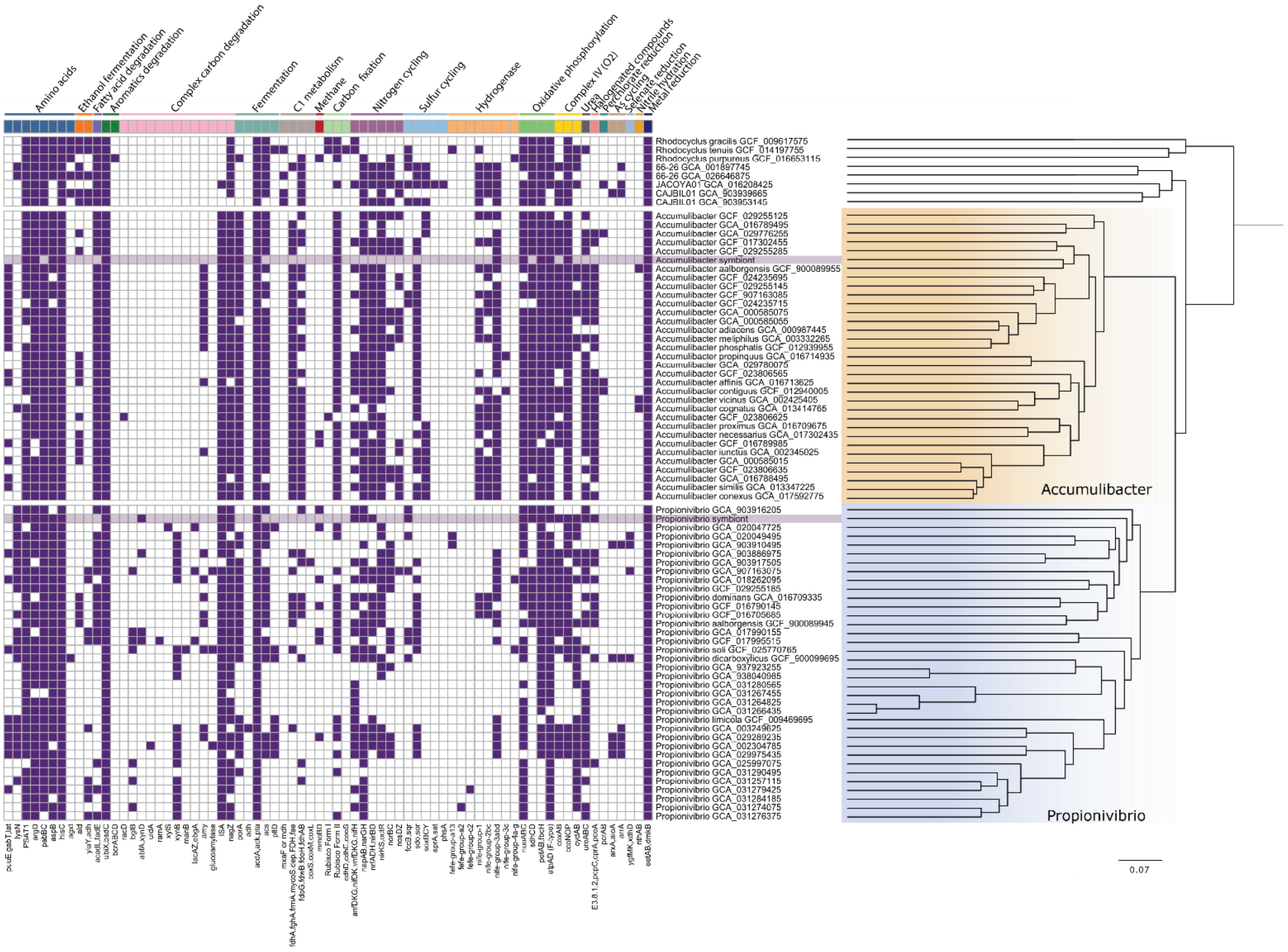

Supplementary Figure 8

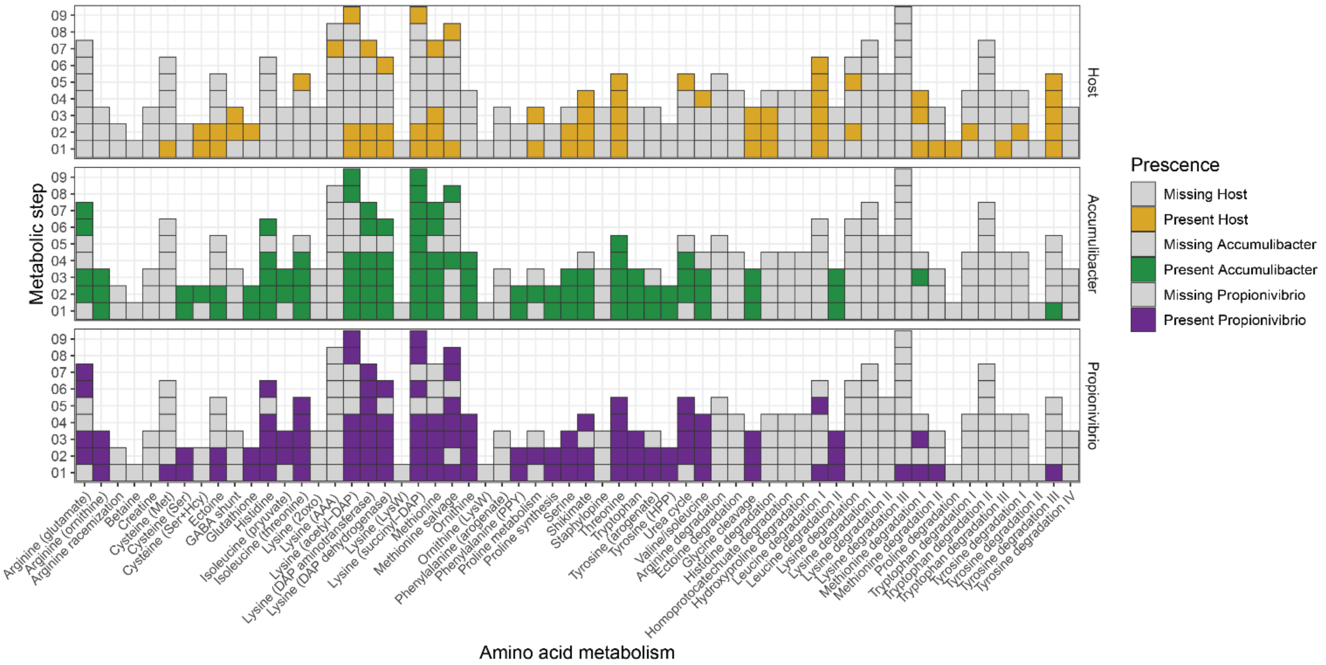

### Supplementary Figure 9

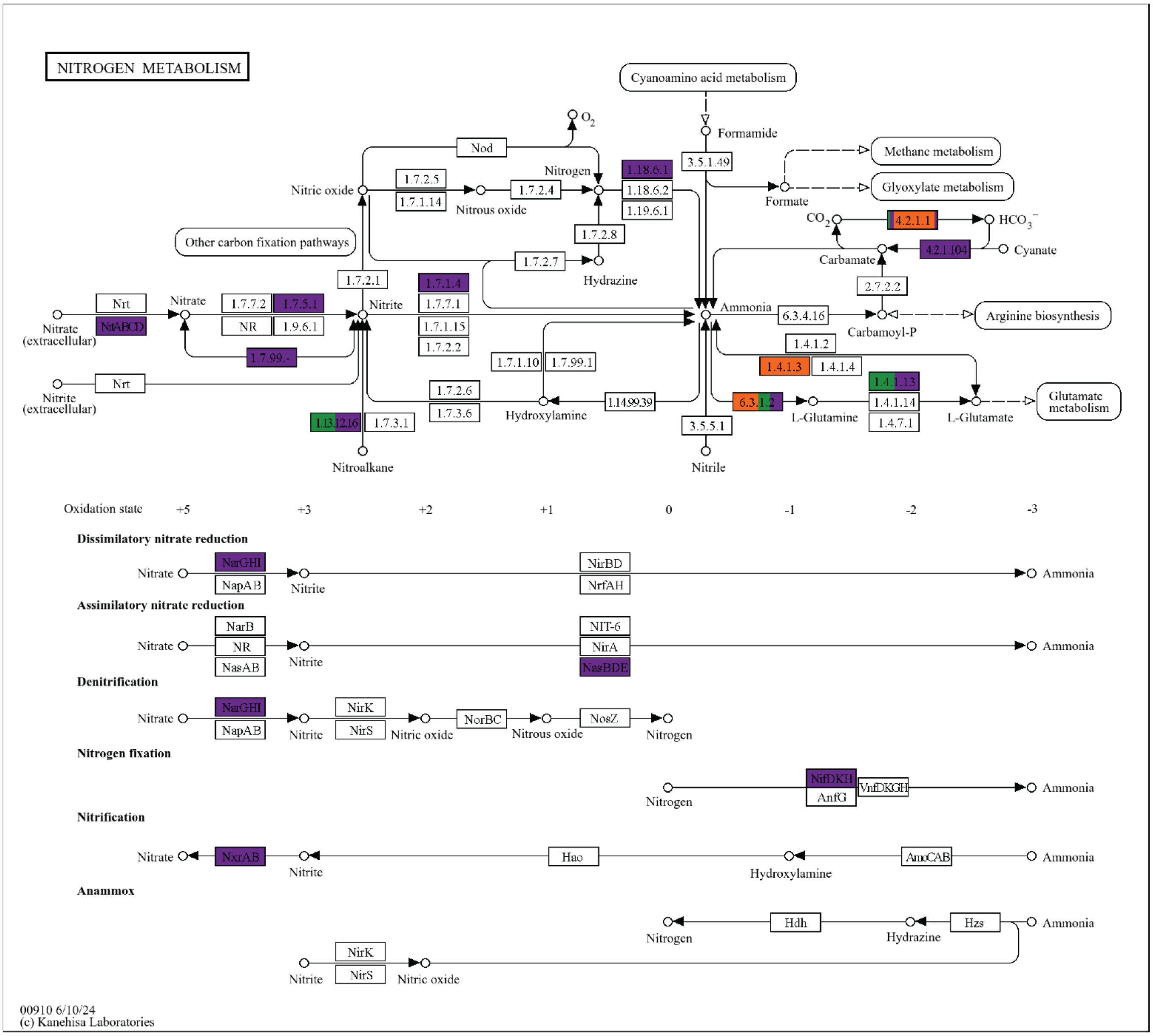

### Supplementary Figure 10

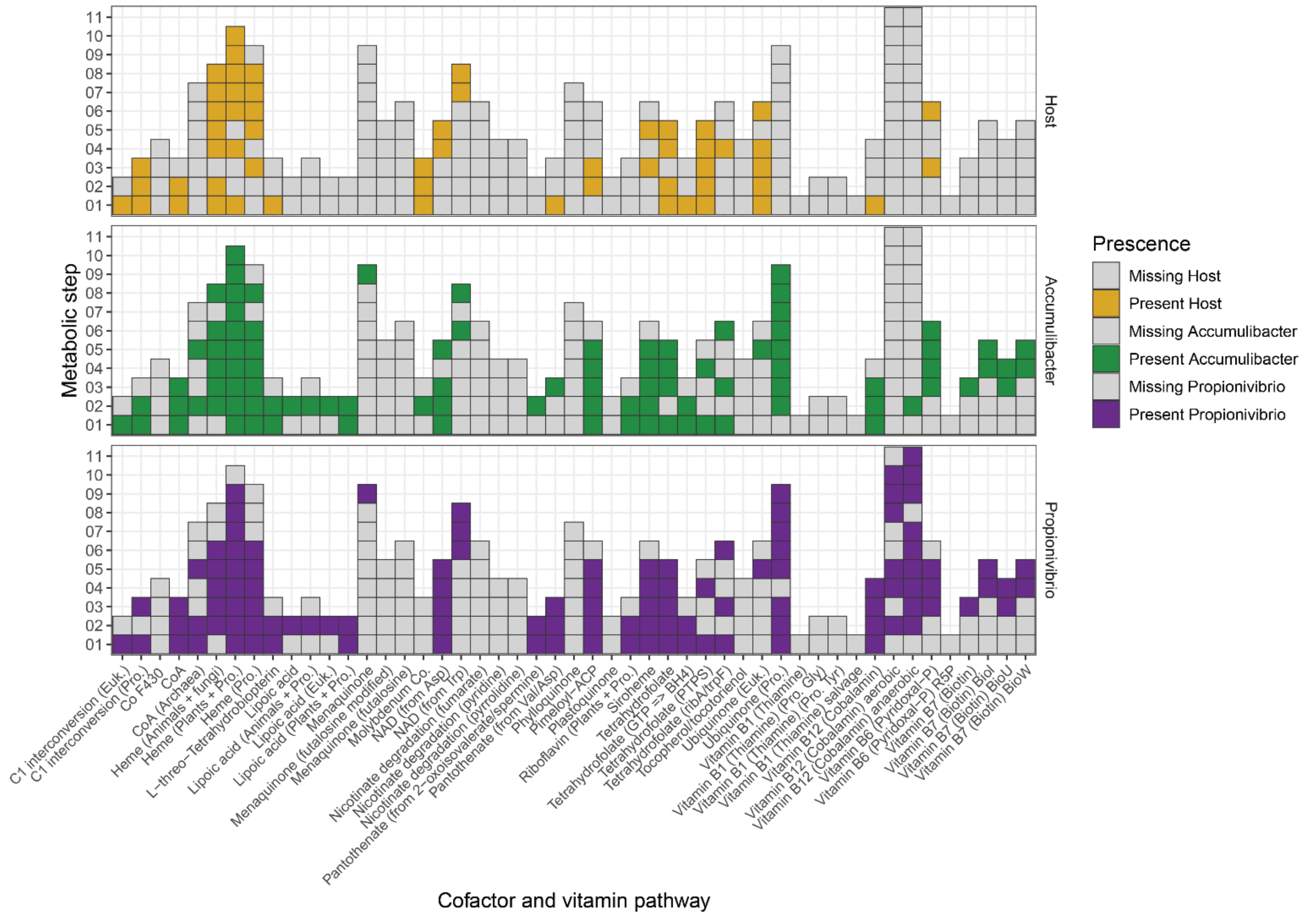

### Supplementary Figure 11

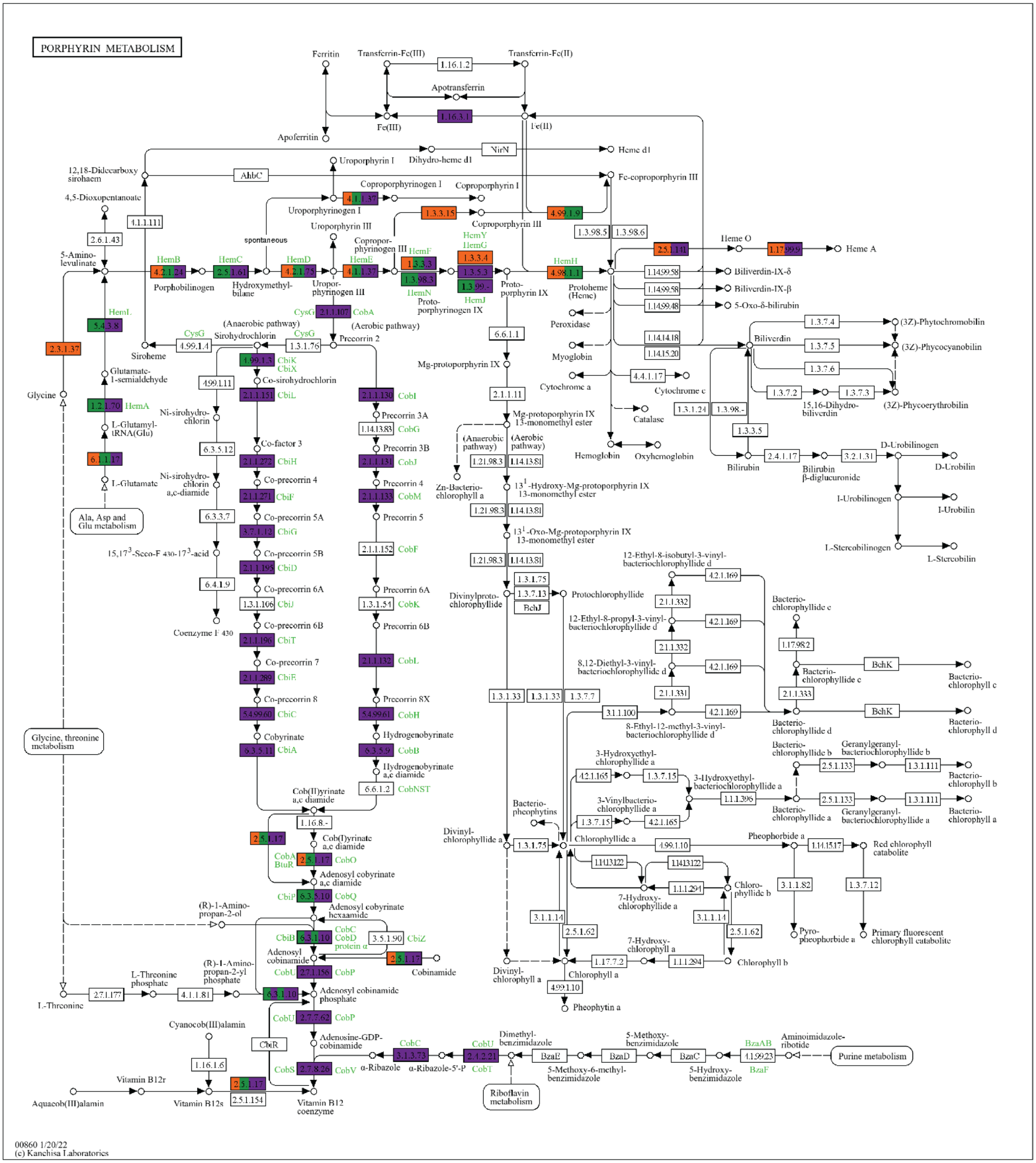

Supplementary Figure 12

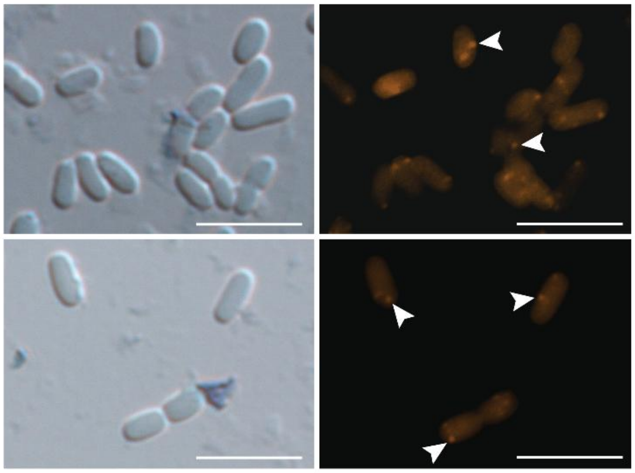

Supplementary Figure 13

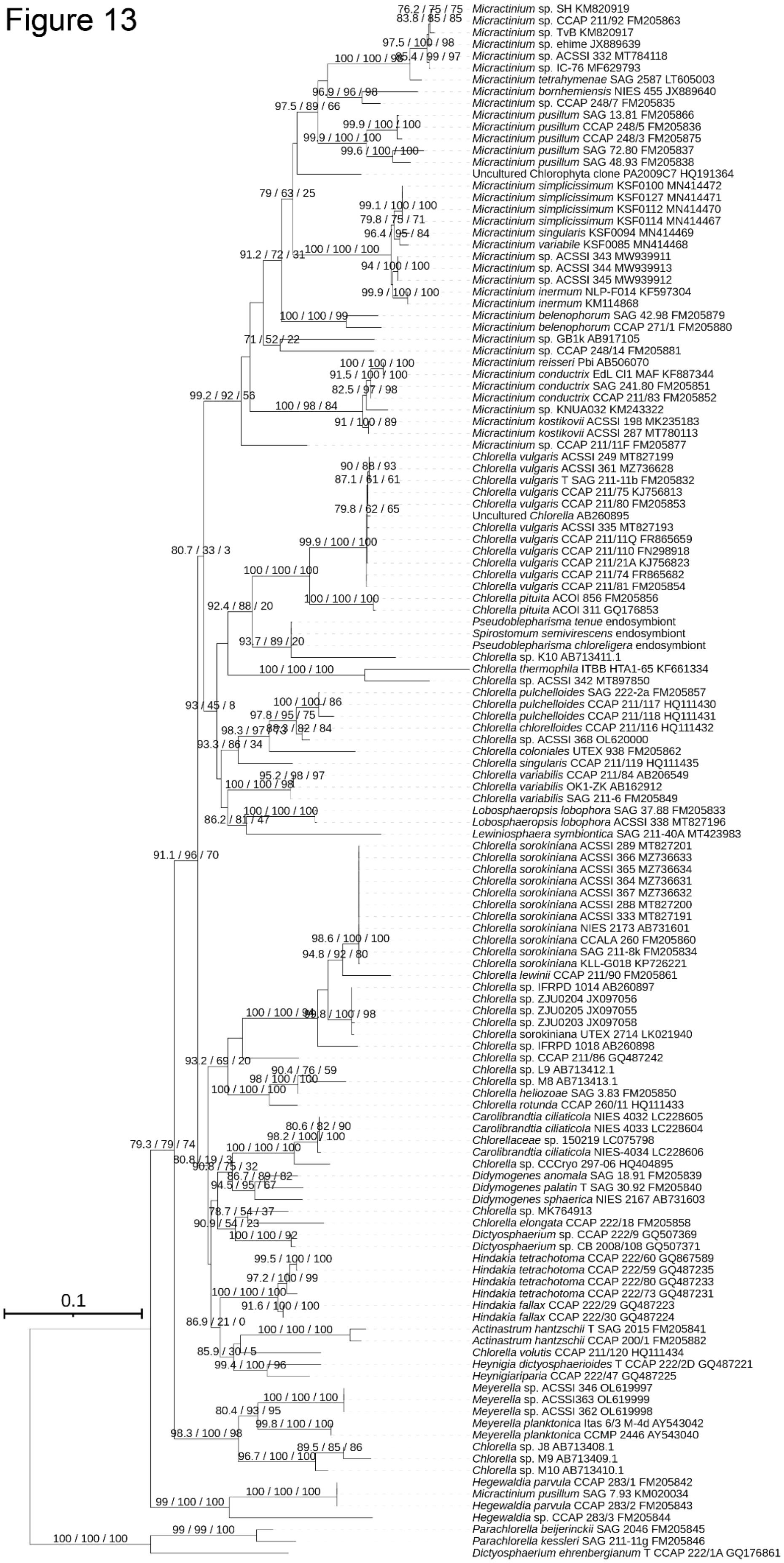

### Supplementary Figure 14

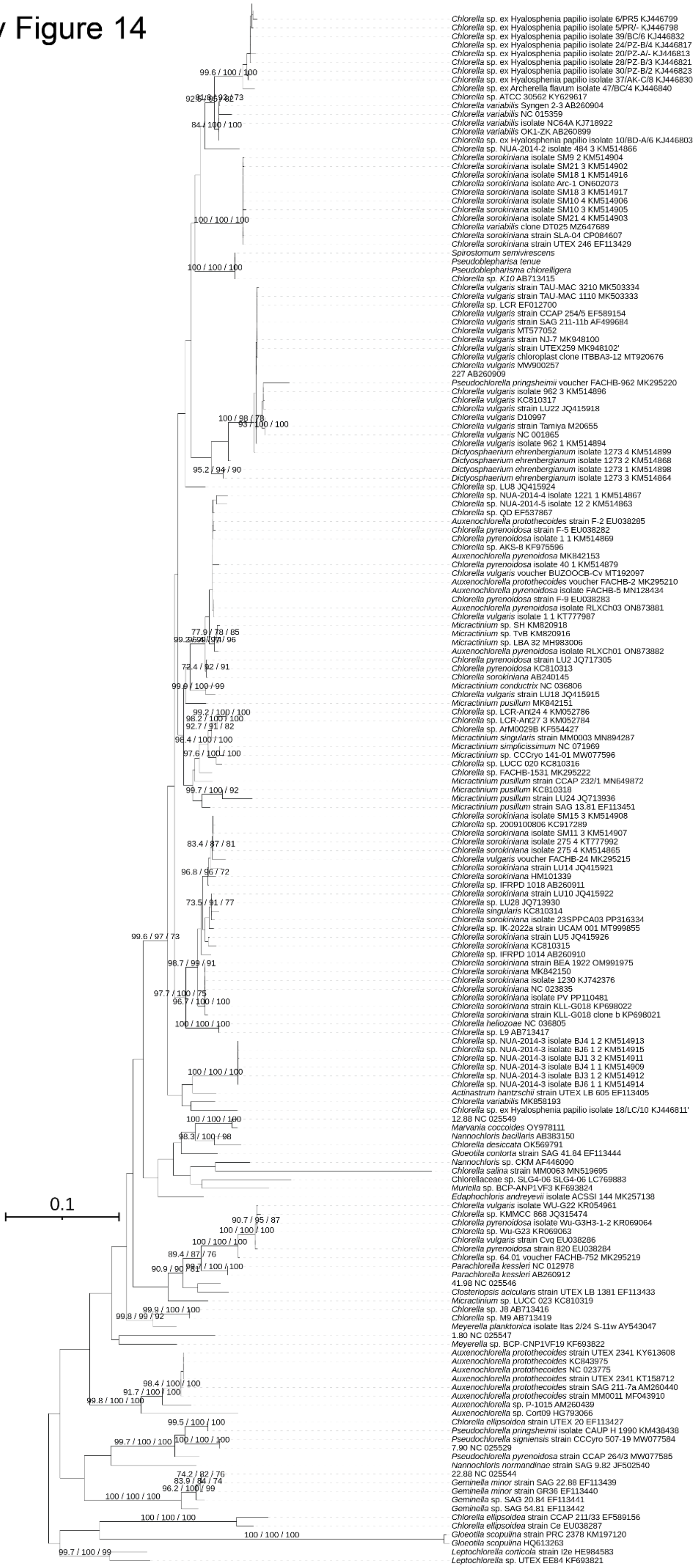

Supplementary Figure 15

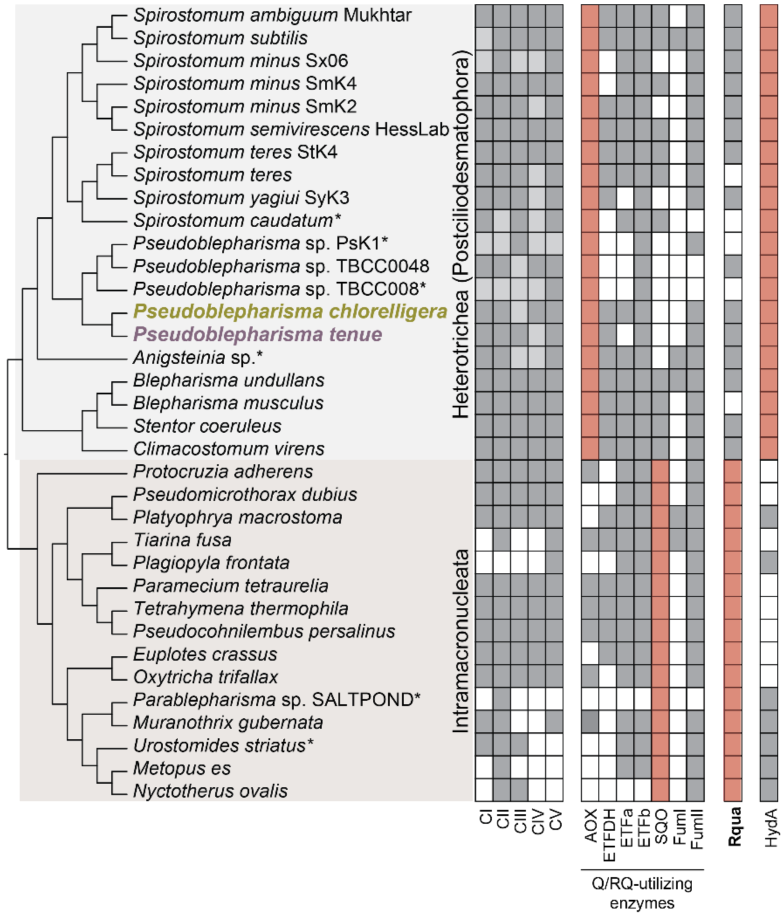
